# Functional specialisation across the first five years of life: a longitudinal characterisation of social perception with fNIRS

**DOI:** 10.1101/2025.07.28.667189

**Authors:** Johann Benerradi, Chiara Bulgarelli, Ebrima Mbye, Borja Blanco, Ebou Touray, Sam McCann, Bosiljka Milosavljevic, Sophie E. Moore, Clare E. Elwell, Anna Blasi, Sarah Lloyd-Fox

## Abstract

Research in typically developing infants has shown robust and consistent brain activation to social versus non-social visual and auditory stimuli in a network of brain regions, including the inferior frontal, anterior temporal and posterior superior temporal cortex. However, large-scale, longitudinal neuroimaging studies across early childhood, particularly in low- and middle-income countries are rare, yet important, given that they offer a powerful means of capturing within-person changes in neurodevelopment and identifying targets for intervention and support. Here we investigated brain responses to social perception longitudinally across the first five years of life of participants in The Gambia with functional near-infrared spectroscopy (fNIRS) within the Brain Imaging for Global Health (BRIGHT) Project. We found that social visual stimuli elicited a selective right posterior temporal response across all six age points studied here (*5*, *8*, *12*, *18*, *24 months* and *3-5 years* of age), with concurrent selective left responses at four of the six time points (*5*, *8*, *18 months* and *3-5 years* of age). Inferior frontal regions showed brain activation at multiple ages, with the youngest (5 *months*) and oldest (*24 months* to *3-5 years*) time points evidencing more widespread frontal social visual responses. Conversely, auditory social stimuli elicited a significantly stronger and more prolonged response than auditory non-social stimuli across all six age points from *5 months* to *3-5 years* across a range of frontal and temporal areas. Finally, from individual infant trajectories of brain activation, we identified different profiles of age-dependent specialisation to auditory social stimuli, with some specialising earlier than others. Future work could take advantage of such profiles to investigate the effects of adversity and resilience factors on neurodevelopment.

## 1. Introduction

Nearly 1 billion children worldwide are growing up in poverty, facing challenges that impact lifelong health and development (“Child Poverty.” UNICEF. https://www.unicef.org/social-policy/child-poverty). These environments, characterised by limited access to essential resources such as adequate nutrition, healthcare, and education, expose infants to various genetic, environmental and psychosocial risks that can have lasting effects on their cognitive and physical development. Exposure to risks such as chronic stress, malnutrition, and inadequate stimulation can disrupt a child’s healthy neural development and affect early developmental trajectories in language and cognition (Jensen et al., 2017, 2019). Indeed, one-third of pre-school-aged children in developing countries fail to reach their developmental milestones in cognitive or socioemotional growth, with the largest number of affected children in sub-Saharan Africa (McCoy et al., 2016). Not only can this lead to deficits in the acquisition of key skills such as delays in attention and communication milestones, it can also lead to poorer academic achievements and a lasting impact into adulthood.

The first 1,000 days of life, between conception and two years of age, are characterised by rapid physical, physiological and psychological development. Investigating development from the earliest stages of life is essential for identifying potential deficits and informing the design of targeted, developmentally appropriate interventions, where needed. Infants’ brain plasticity and specialisation in response to environmental factors plays a critical role in early development. Mapping brain function across the first two years of life is key to understanding the ages at which detrimental effects appear, and the contextual factors most closely associated with subsequent neural or cognitive differences up to five years when children reach school age.

A key area of early development is how infants respond, interpret and learn from their social world. The capacity to engage and communicate in a social world is one of the defining characteristics of the human species. Given that human infants are reliant on their caregivers for an extensive period of their development, opportunities for early interactions with their social environment are largely driven by their caregivers. Therefore, risk factors that disrupt caregivers’ lives (i.e. unstable family income, food insecurity, mental health issues, life stressors, climate impacts) will also impact their ability to provide sensitive, responsive and extensive opportunities for social interaction, which in turn may impact early neural trajectories of our capacity to perceive differing social cues. The link between life stressors, early adversity, socioemotional development and social cognition has been widely studied over the last century. Previous research following the Great Depression of the United States of America found that economic loss led to changes in the quality of caregiving which in turn associated with early socioemotional development (Elder et al., 1979). Grantham-McGregor and colleagues (Grantham-McGregor et al., 1991) investigated the effects of nutritional supplementation and psychosocial stimulation for infants living in low resource contexts in Jamaica, finding that those infants with caregivers who learnt more sensitive caregiving practices and social interaction skills had enhanced cognitive development at pre-school age, and positive benefits extending into adulthood (Walker et al., 2022). Furthermore, predictive modelling across 35 low- and middle-income countries (LMICs) of the Early Childhood Development Index (ECDI) has shown that socio-emotional development of pre-schoolers (3-4 years) is impacted more extensively (26.2% of cohort with low scores) than cognitive development (14.6%) by associated risk factors (McCoy et al., 2016).

Recent efforts to understand the neural mechanisms by which early social cognition and socioemotional development are impacted by adversity and risk factors are limited but provide some insight. In a pilot work conducted in The Gambia using functional near-infrared spectroscopy (fNIRS) to study social perception neural responses, we found evidence of age-related changes in visual- and auditory-social selectivity across the first two years of life (Lloyd-Fox et al., 2013; 2017). However, this research had some limitations: (i) while the study was based in a low resource context, it did not directly study associations with adversity factors due to the low sample size; (ii) the study investigated responses at 1 and 24 months cross-sectionally, or with a more limited longitudinal sample from 9-16 months; and finally, (iii) the neuroimaging headgear was a small feasibility kit that only covered one region of interest in the right hemisphere making the investigation of broader social network activation impossible. On the other hand, a study in Bangladesh has shown that brain activation to social compared to non-social stimuli in 6-, 24- and 36-month-olds was related to poverty, maternal education, maternal stress, caregiver violence and verbal abuse (Perdue et al., 2019; Pirazzoli et al., 2022). While this study contributed considerably to our understanding of the neural mechanisms of social cognition and associations with adverse contextual factors, it studied each participant at only 1-2 time points, with the 6-24-month and 36-month cohorts being separate. An objective of the work presented herewith will therefore be to investigate changes in social selectivity with a larger individual age span than previous work to validate findings which described a switch from non-vocal selectivity to vocal selectivity during the first year of life (Lloyd-Fox et al., 2012; 2017; Grossmann et al., 2010).

Conducting longitudinal studies across an extensive early developmental window (e.g. from birth to school age) is essential, as it allows researchers not only to identify when individual skills and neural markers emerge, but also to map the trajectories of cognitive development over time. When combined with other contextual measures, this approach is important for investigating the factors that influence developmental trends and for identifying distinct developmental profiles. Longitudinal studies offer a much larger window to capture age-related changes that may develop rapidly at key developmental stages, while also enabling a clearer understanding of causal relationships.

However, longitudinal neuroimaging studies are scarce (Azhari et al., 2020), and very few neurodevelopmental studies have been conducted in LMICs (Blasi et al., 2019), with literature identified from just two contexts: Bangladesh (Perdue et al., 2019) and India (Wijeakumar et al., 2019). The scarcity of neuroimaging studies in LMICs is partly due to the typically high cost of neuroimaging equipment and shortage of well-resourced research labs (which until recently was the only context in which neuroimaging research studies were routinely undertaken). However, recently, the adoption of new more affordable, flexible and easy-to-use neuroimaging technologies such as fNIRS and electroencephalography (EEG) in LMICs has made it possible to measure functional brain responses in more varied contexts (Blasi et al., 2019; Gervain et al., 2023). In 2014, the interdisciplinary, longitudinal Brain Imaging for Global Health (BRIGHT) Project was launched with the aim to expand diversity in neurodevelopmental studies and establish brain function-for-age curves in infants from high-(UK) and low-resource (The Gambia) settings in order to gain an insight into the effects that contextual factors related to living in a low-resource context may have on infant development (Lloyd-Fox et al., 2024). To achieve this aim, a wide range of measures were employed, including neuroimaging (fNIRS and EEG), eye tracking and physiological measures, alongside behavioural and cognitive assessments and measures of quality of the child’s environment and caregiving practices. These were complemented by a wealth of data on both biological (physical size and biomarkers of nutritional status and inflammation) and psychosocial risk factors (e.g., questionnaire-based measures of adversity exposure and parental mental health).

To investigate the developmental trajectory of our capacity to perceive social cues across early childhood, the BRIGHT Project utilised a social perception fNIRS task to capture visual and auditory social neural responses from birth to 5 years of age. To our knowledge this is the first neuroimaging study to measure developmental longitudinal neural trajectories of social perception across this number of time points, with this extensive early childhood age range and with this sample size. Therefore, while in future work we aim to include this data in modelling pathways to investigate the impact of adversity, a first priority was to better understand the origin, specialisation patterns and developmental trajectories of visual and auditory social perception and isolate potential biomarkers.

Due to the sample size and number of time points, this work is also motivated by the aim of better understanding whether methodological and analytical parameters may impact accurate profiling of developmental trajectories. One motivation to explore individual differences further is that previous literature indicates that when comparing speech, non-speech human vocalisations and non-human biological or non-biological sounds, infants display evidence of developmental specialisation to vocal stimuli across the first year of life (Lloyd-Fox et al., 2012; 2017; Grossmann et al., 2010; Minagawa-Kawai et al, 2011; Shultz et al., 2014). While this has been validated through several replication studies, in different populations and with the use of varying stimuli, some differences have arisen. This may suggest considerable individual infant variability of specialisation, which may be related to contextual factors of interest. However, it is possible that the reported significant findings may also vary for methodological reasons such as:

- which brain region was investigated (i.e. temporal only, left hemisphere vs bilateral, several social or language network regions);
- how the analyses were conducted (i.e. selection of a specific time windows of interest post stimulus onset used for feature extraction, pre-defined regions of interest from previous literature, hardware constraints on extent of the anatomical region that can be investigated);
- which stimulus choice of comparison was used (i.e. speech vs communicative vocal sounds (yawn or laugh) vs non-speech biological sounds (walking), or vocal (coughing, crying, laughing or yawning) vs non-vocal sounds (water or toys));
- which part of the haemodynamic signal was focused on (i.e. for oxyhaemoglobin (HbO) only or both HbO and deoxyhaemoglobin (HbR), for stimulus comparison, focusing on optimal parameters to capture stimulus of interest rather than baseline);
- which age point the study focused on (i.e. studies tend to be cross-sectional or focus on a narrow age span such as 1 to 4 months or 4 - 7 months).

The larger sample size and the wide longitudinal age range including multiple testing sessions used in the BRIGHT project can allow us to investigate some of these factors. Firstly, we can research age-related changes more comprehensively. Secondly, we can investigate whether the analytical choices bias findings to be significant for the stimulus of interest (i.e. an analytical time window optimised for the social response may undervalue the response seen to the contrasting auditory non-social stimuli). Thirdly, we can explore whether notable age-related group level and individual differences may arise if we widen our scope of potential biomarkers to include a range of haemodynamic response characteristics. These include the standard approach to identify the localised peak response within a given region using an average haemodynamic change analysis, as well as investigating time-to-peak (latency), magnitude of response to stimulus relative to baseline and spatial extent of the significantly activating brain regions (i.e. narrow area of activation vs widespread activation across brain regions).

This paper is divided into three analyses:

1. First, we study the characteristics of the hemodynamic response time courses during the social stimuli at each session from 5 months to 5 years of age.
2. Second, we study the spatial distribution of brain activation during the social task cross-sectionally at each session from 5 months to 5 years at the group-level.
3. Third, we extract participants’ longitudinal trajectories from 5 months to 5 years, creating individual profiles to explore the neural bases underlying the differences between participants’ social specialisation.

## 2. Methods

### 2.1 Participants

Participants for this study were recruited as part of the BRIGHT Project (Lloyd-Fox et al., 2024), from villages across the rural West Kiang District in The Gambia, where a field station of the MRC Unit The Gambia at the London School of Hygiene and Tropical Medicine is based. Ethnicity in the West Kiang District of The Gambia is predominantly Mandinka (79.9 % of the population (Hennig et al., 2017)) with its unique language and cultural characteristics. The local population commonly experience seasonal food insecurity, and infections and other health challenges are common, especially among infants and young children, often leading to growth faltering (Nabwera et al., 2017; Schoenbuchner et al., 2019).

Demographics from the full Gambian cohort in the BRIGHT Project can be found in Lloyd-Fox et al., 2024: in summary, families live in multigenerational households with up to 36 members per compound. Polygamy was common within the cohort with 38.8 % of fathers having more than one wife. Consequently, while mothers had on average 4.4 children, including the infant enrolled in the study, fathers had on average 6.9, with a range of 1-23 children attributed to a single father. For the generation of parents within our cohort, formal schooling was not readily available when they themselves were children, therefore, on average, mothers and fathers within the study had completed three and four years of schooling respectively. To avoid confounds caused by multiple translations of the study protocols, only families where Mandinka was spoken as the first language were recruited into the study. In addition, 16.7 % of primary caregivers reported that they spoke a second language and 3.5 % of primary caregivers spoke three languages.

Participating families were recruited into BRIGHT antenatally, were seen at home for a neonatal examination in the immediate postpartum period and were then invited to attend scheduled assessments at 7 timepoints post-partum (*7-14 days* (at home), *1*, *5*, *8*, *12*, *18* and *24 months* of infant age). The same cohort was then followed up at a single further timepoint when they were aged between 3 and 5 years of age (the BRIGHT Kids study). 204 infants were initially enrolled at *1 month* of age.

Ethics approval was obtained from the joint Gambia Government - MRC Unit The Gambia Ethics Committee (SCC 1351 for BRIGHT and Project reference 22737 for BRIGHT Kids), and informed consent was obtained from carers/parents in writing or via thumbprint if they were unable to write.

### 2.2 Experimental design

The auditory-visual social perception paradigm used for this study was first described by Lloyd-Fox and colleagues (Lloyd-Fox et al., 2014). This was the first paradigm presented to participants as part of a battery of fNIRS tasks (Lloyd-Fox et al., 2024) implemented at the *5*, *8*, *12*, *18*, *24 months*, and *3-5 years* study visits. fNIRS data for the social perception task were collected from 140 infants at *5 months* of age; 126 at *8 months*; 132 at *12 months*; 121 at *18 months*; 130 at *24 months*; and 163 at *3-5 years*. The task was also presented at *1 month*, but that session was not included here as the paradigm was different, with infants being asleep and no *visual social (silent)* condition.

The trials were presented with a period of baseline between each. The sequence of stimulus presentation has been used in previous research (Lloyd-Fox et al., 2012; Lloyd-Fox et al., 2013; Blanco et al., 2023) and is presented in Figure 1.

**Figure 1.**
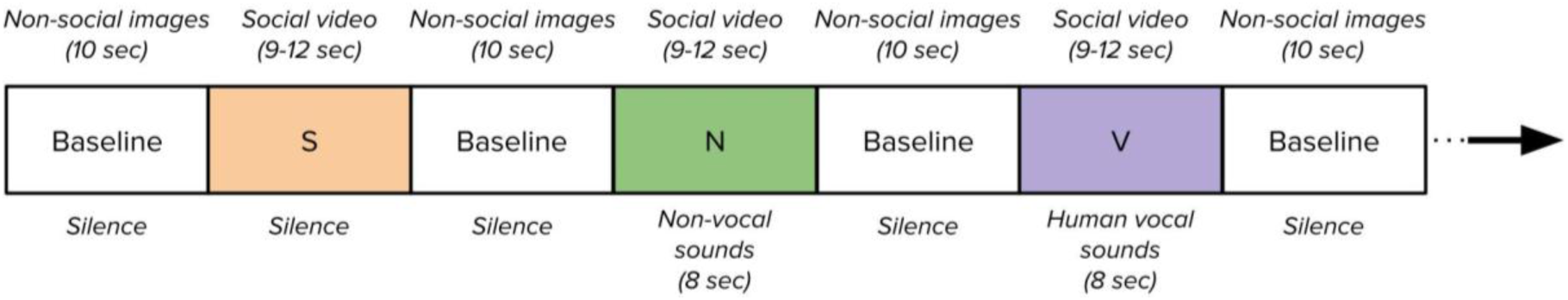
Illustration of the social task experiment paradigm showing the sequence of trials with duration of the stimuli (S: visual social (silent); N: auditory non-social (non-vocal); V: auditory social (vocal)).

Three conditions (*visual social (silent)*, *auditory social (vocal)*, *auditory non-social (non-vocal)*) were presented in the same pseudo-randomised order across infants in a repeating loop (*visual social (silent)* → *auditory non-social (non-vocal)* → *auditory social (vocal)* → *visual social (silent)* → *auditory social (vocal)* → *auditory non-social (non-vocal)* → …) of trials until the infants became bored or fussy as judged by the experimenter who was monitoring their behaviour, up to a maximum of 8 trials per condition. During each experimental trial, a silent social video of 9 to 12 seconds with a local actor performing nursery rhyme gestures was played. These consisted of full-colour, life-size (head and shoulders only) social videos of adults (Gambian nationals) who either moved their eyes left or right or performed hand games: “Peek-a-boo” and “Incy-Wincy Spider”. To ensure infants’ continuous attention, especially since the social visual stimuli was also presented during auditory trials, there were six different visual social videos (two actors; three types of social video). These visual stimuli were displayed on a 24-inch plasma screen with a viewing distance of approximately 1 meter. For the *auditory social (vocal)* and *auditory non-social (non-vocal)* trials, auditory cues were also played at the onset of the trials with a duration of 8 seconds. The content and duration of the social videos were not synchronised with the auditory stimuli. The 8-second auditory stimuli consisted of four different sounds (of vocal or non-vocal stimuli) presented for 0.37–2.92 seconds each, interleaved by short silence periods (of 0.16–0.24 seconds). The two auditory conditions were equivalent in terms of average sound intensity and duration (p-value = 0.65). In the *auditory social (vocal)* condition, the video was accompanied by non-speech vocalisations (sequence of coughing, crying, laughing and yawning). In the *auditory non-social (non-vocal)* condition, it was environmental sounds (that were not human or animal produced) familiar to the participants (sequence of running water, bells, and rattles). In the *visual social (silent)* condition, the video was not accompanied by any sounds. A restriction of studying auditory processing in awake infants of this age is that they need to be presented with concurrent visual stimulation to reduce infant movement and thus artefact in the signal. Therefore, in line with previous work using these stimuli, we chose to employ the same visual stimuli during the presentation of the auditory stimuli as the one used when auditory stimulation was absent. Each experimental condition was separated by a baseline of 10 seconds with a pseudo-random sequence of images of objects, amenities or environment familiar to the participants, each image presented for 1-3 seconds.

Videos of the participants were recorded during the task, and participants’ gaze was monitored using an eye tracker (Tobii TX-300, Tobii AB, Sweden). This enabled the identification of trials during which the participants did not pay attention to the task. Trials with less than 60 % engagement were discarded from the analysis.

### 2.3 Data acquisition

fNIRS data were collected using a continuous wave NTS fNIRS System (Gowerlabs Ltd. London, UK) with two wavelengths of 780 and 850 nm, and a sampling frequency of 10 Hz (Everdell et al., 2005). The fNIRS sensor included 12 sources and 14 detectors for a total of 34 channels located over the frontal and temporal brain regions bilaterally for participants aged between *5 months* and *3-5 years* (Figure 2). At *3-5 years*, new optical fibres were used, and the custom-made silicone band holding the optode arrays was replaced by a modified EasyCap (EASYCAP GmbH), which facilitated replicability of headgear positioning with the older participants who had more hair than young infants.

**Figure 2.**
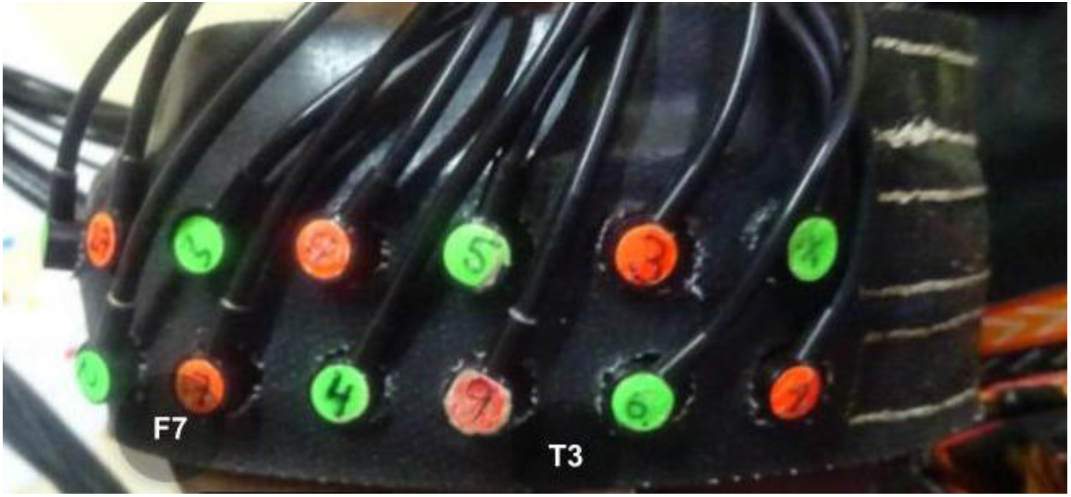
NTS fNIRS System used for the BRIGHT study with reference 10-20 positions for participants aged between 5 months to 3-5 years (12 months example on the photo).

Before starting the session, infants’ head measurements were taken (head circumference, ear-to-ear measurements over the forehead and over the top of the head) to guide alignment of the headgear with the 10-20 system anatomical landmarks (pre-auricular points and nasion) (Lloyd-Fox et al., 2014). Once the headgear was securely placed on the participant’s head, photographs of the infants wearing the headgear were taken. Whenever possible, photographs were taken at the end as well. Pictures were used to assess the headgear placement, enabling us to reject recordings for which the displacement of reference optodes was greater than 1.6 cm based on the known inter-optode distance (Blasi et al., 2014). The array was registered onto age-appropriate brain templates with anatomical labels (Collins-Jones et al., 2021). This co-registration was completed for all time points except the *3-5 years* age group. However, as after a phase of rapid growth of the brain in infancy (reaching 80 % of the adult size at *24 months*) the rate of change of brain structure slows (Gilmore et al., 2018), we assumed that channel mapping would not change greatly between *24 months* and *3-5 years*. Consistent with this, head circumference at *24 months* (46.58 ± 1.47 cm) measured about 95 % of the head circumference at *3-5 years* (49.20 ± 1.43 cm) (see Table 1). The cortical areas that the channels covered for infants aged *5 months* to *3-5 years* ranged from the inferior frontal gyrus at the front of the array, to the posterior sections of the superior, middle, and inferior temporal gyri at the back of the array.

**Table 1.** Participant information describing for each session the number of participants with data analysed (N), the percentage of male and female participants, the mean age ± standard deviation (SD) at the session calculated from the date of birth (in months), the mean ± SD weight-for-height Z-score (WHZ), the mean ± SD head circumference (HC, in cm), the mean ± SD number of retained channels (out of a total of 34), and the mean ± SD number of retained trials per condition (out of a total of 8 maximum).

| Session | N | Male/Female | Age (months) | WHZ | HC (cm) | Number of channels | Trials per condition |
| --- | --- | --- | --- | --- | --- | --- | --- |
| 5 mo | 127 | 48%/52% | 5.26 ± 0.30 | -0.19 ± 1.01 | 41.28 ± 1.25 | 30.61 ± 3.43 | 5.55 ± 0.92 |
| 8 mo | 110 | 50%/50% | 8.11 ± 0.37 | -0.37 ± 1.03 | 42.85 ± 1.34 | 30.79 ± 3.06 | 5.14 ± 0.83 |
| 12 mo | 116 | 49%/51% | 12.23 ± 0.45 | -0.57 ± 1.02 | 44.39 ± 1.37 | 30.56 ± 2.81 | 5.20 ± 0.93 |
| 18 mo | 111 | 52%/48% | 18.34 ± 0.60 | -0.60 ± 0.93 | 45.91 ± 1.48 | 29.70 ± 3.46 | 5.14 ± 0.86 |
| 24 mo | 112 | 46%/54% | 24.39 ± 0.77 | -0.42 ± 0.94 | 46.58 ± 1.47 | 28.71 ± 3.68 | 5.38 ± 0.66 |
| 3-5 years | 124 | 56%/44% | 48.98 ± 6.92 | -0.78 ± 0.93 | 49.20 ± 1.43 | 32.26 ± 2.86 | 5.67 ± 0.44 |

### 2.4 fNIRS data pre-processing and quality control

The data processing followed the pipeline described by Bulgarelli and colleagues (Bulgarelli et al., 2020) and was implemented in *MATLAB* using *Homer2* (Huppert et al., 2009). First, channels with very low intensity (lower than 3.10^−4^, specific to the NTS fNIRS system) or low signal quality assessed using *QT-NIRS* (SCI = 0.7, cardiac power = 0.1 µV) (Pollonini et al., 2014; Hernandez and Pollonini, 2020) were discarded. Light intensities were converted into optical densities and then motion artefacts were detected using *Homer2* to be corrected using a combination of Spline interpolation (p=0.99) and wavelet filtering (iqr=0.8) as previously suggested (Di Lorenzo et al., 2019). Trials that still showed evidence of motion artefacts after correction were then rejected. Moreover, trials that were attended for less than 60 % by the infants were also rejected. Data were bandpass filtered (0.02-0.6 Hz) in order to remove instrumental and systemic noise while keeping frequencies related to the hemodynamic response function (HRF) and experiment paradigm (around 0.1 Hz). We then converted the signal into HbO and HbR concentration changes using the modified Beer-Lambert law (Delpy et al., 1988) and baseline corrected using the pre-trial baseline of 2 seconds prior to the trial trigger onset. Lastly, concentration changes were block averaged up to 20 seconds post-stimulus onset and baseline corrected using 2 seconds prior to the stimulus onset. The block averaging method was preferred here compared to the GLM, given its simplicity and the evidence that both approaches eventually lead to similar conclusions (Luke et al., 2021). Participants were excluded from data analyses at a specific session if they had fewer than 3 remaining trials per condition or less than 60 % valid channels after preprocessing and looking time analyses.

### 2.5 *First analysis*: profiling the fNIRS responses

Statistical analyses were performed using a combination of in-house Python 3 scripts based on *SciPy* (Virtanen et al., 2020) and *statsmodels* (Seabold and Perktold, 2010).

Firstly, we examined the characteristics of the haemodynamic response to the stimuli by block averaging each infant’s haemodynamic responses to the different conditions (*visual social (silent)*, *auditory non-social (non-vocal)*, *auditory social (vocal)*) within each session. We explored whether we would observe age-related and stimulus-related differences in haemodynamic profiles. To investigate whether the magnitude of the haemodynamic response to the presentation of the experimental condition trials (relative to the previous baseline condition response) was influenced by the age point and stimulus type, we extracted the average maximum amplitudes for HbO and average minimum amplitudes for HbR for each infant and ran two-way ANOVA tests. To investigate the latency of the peak response, we examined the time at which the peak (maximum HbO or minimum HbR) of the haemodynamic response profile was observed for each stimulus type and age point and tested significance again with two-way ANOVA tests.

### 2.6 *Second analysis*: cross-sectional group level analysis

Following up the profiling of fNIRS HRFs, we investigated brain responses to social stimuli cross-sectionally at the group level. This was done using the mean HbO and HbR concentration changes over a 4-second range around the average time-to-peak (i.e. HbO time-to-peak was determined by averaging all trials and channels within a given condition at the group level, also averaged across the two compared conditions in the case of a contrast). A 4-second span was selected to include a sufficient individual variability of time-to-peak around the average time-to-peak across participants. The choice of extracting the mean around the expected peak and not on a fixed time window was motivated by limitations from previous work. Indeed, in our previous feasibility research using the same visual and auditory stimuli within this rural Gambian population we found preliminary evidence of differences in the timing and shape of the hemodynamic response, which potentially also differed between social and non-social conditions and age of participant (Lloyd-Fox et al., 2017). Therefore, our approach decreased the likelihood of preferentially biasing one condition over another, or one age point over another.

One-way channel-by-channel comparisons were performed using t-tests to assess brain activation during the *visual social (silent)* condition (one-sample t-tests, to investigate the response to *visual social (silent)* relative to the final 2 seconds of the previous non-social baseline trial) or the contrast between *auditory social (vocal)* and *auditory non-social (non-vocal)* conditions (paired t-tests). Activation was defined as a statistically significant increase in HbO and/or statistically significant decrease in HbR with a p-value smaller than 0.05. False discovery rate (FDR) correction (Benjamini and Hochberg, 1995) was performed at the channel level (34 tests, one per channel), for each chromophore separately. For the auditory contrast more specifically, we defined a selectivity towards *auditory social (vocal)* as a significant activation for *auditory social (vocal)* coupled with a greater response to *auditory social (vocal)* than *auditory non-social (non-vocal)*.

### 2.7 *Third analysis*: longitudinal analysis of individual cortical specialisation

#### Feature extraction

Following the cross-sectional analysis, we aimed to investigate age-related changes in auditory social selectivity, with the comparison of the *auditory social (vocal)* and *auditory non-social (non-vocal)* stimuli at the participant level. Here, we focused on HbO as a biomarker of social selectivity: this was driven by the fact that it has a higher contrast-to-noise ratio than HbR (Strangman et al., 2002), especially with infants (Cristia et al., 2013), which therefore makes it a more suited feature to use at the individual level. This choice was further backed by our analysis of the fNIRS responses which confirmed that task responses were positive in HbO and negative in HbR (Figure 6), without signs of major motion artefacts. Based on co-registration (Collins-Jones et al., 2021) for each hemisphere, the optode array was divided into three regions of interest from anterior to posterior (see Appendix 1): the *frontal* region of interest was composed of the middle/inferior frontal gyri; the *anterior temporal* region of interest was composed of the anterior temporal gyri (up to the T4/T3 10-20 landmark); the *posterior temporal* region of interest was composed of the posterior temporal gyri (from the T4/T3 10-20 landmark and back). This enabled us to average channels by region of interest in order to study comparable anatomical brain regions across ages, accounting for head growth and head shape changes. Each region of interest ranged from 4 to 6 channels, specific enough to investigate previously identified regions of interest from literature (Pernet et al., 2015), but with enough channels to mitigate missing channels and the use of fMRI templates as opposed to fMRI scans from each participant. Selectivity towards social stimuli was defined as an anterior temporal HbO response which was positive for the *auditory social (vocal)* condition and greater for the *auditory social (vocal)* than *auditory non-social (non-vocal)*. Selectivity towards non-social stimuli was defined as an anterior temporal HbO response which was positive for the *auditory non-social (non-vocal)* condition and greater for the *auditory non-social (non-vocal)* than *auditory social (vocal)*. The remaining participants were defined as non-selective towards auditory social (vocal) nor auditory non-social (non-vocal).

#### Individual social selectivity responses across ages

Using the extracted categorical features, we described the evolution of the proportion of participants showing a social selectivity across ages, and used a Bayesian Generalized Linear Mixed Model (GLMM) with a binomial link function to analyse the relationship between age and the social selectivity binary outcome (auditory social selectivity or not). The model included random intercepts for participants to account for within-subject variability. Then, to describe the social selectivity responses further, we investigated the evolution of social selectivity responses at the individual level for participants with valid data at each session during the first year of life (*5*, *8* and *12 months,* N=50) in each of the 3 regions of interest bilaterally.

#### Developmental trajectories

Finally, continuing analysis with the 50 participants with full data over the first year of life, we proposed a way to group participant into social selectivity profiles based on their age of first social selectivity: either first social selectivity at *5 months*, *8 months*, *12 months*, or categorised as a *remaining group* if they did not show social selectivity by then. This enabled us to visualise their brain response trajectories to explain the different social selectivity profiles.

## 3. Results

Overall, the number of participants with valid fNIRS data varied from 110 to 127, depending on the session. The detail of sample retention and attrition is presented in a flowchart in Figure 3. Of those infants who took part in the study at each age point data quality was overall high with retention ranging from 76 to 92%, showing that these paradigms are suitable for use across this age range. Participant characteristics for each session are presented in Table 1.

**Figure 3.**
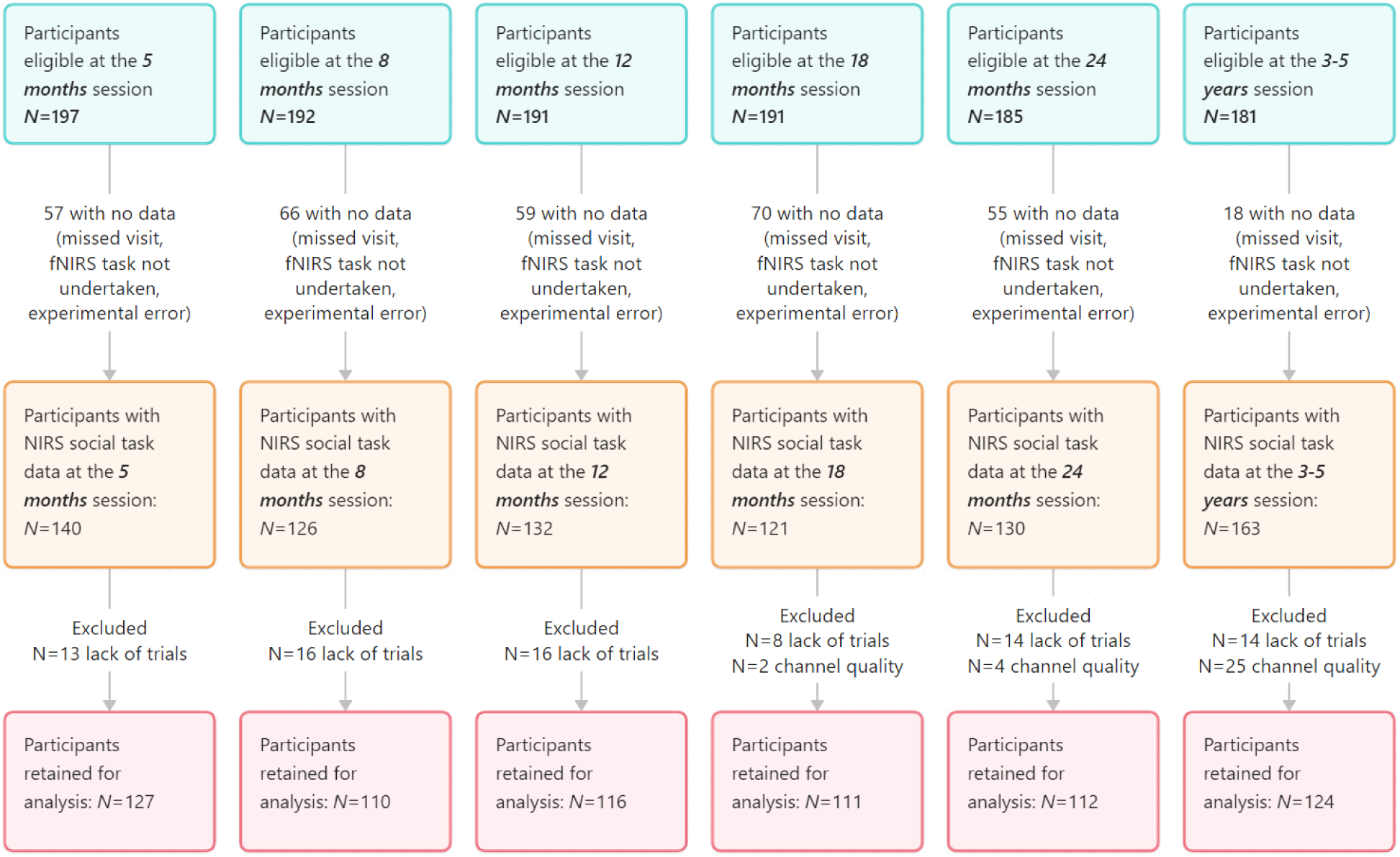
Flowchart of available fNIRS data at each age point.

On average, participants had 3.5 sessions with fNIRS data retained for analysis, with 110 participants having valid fNIRS data on 4 sessions or more between *5 months* and *3-5 years* (Figure 4).

**Figure 4.**
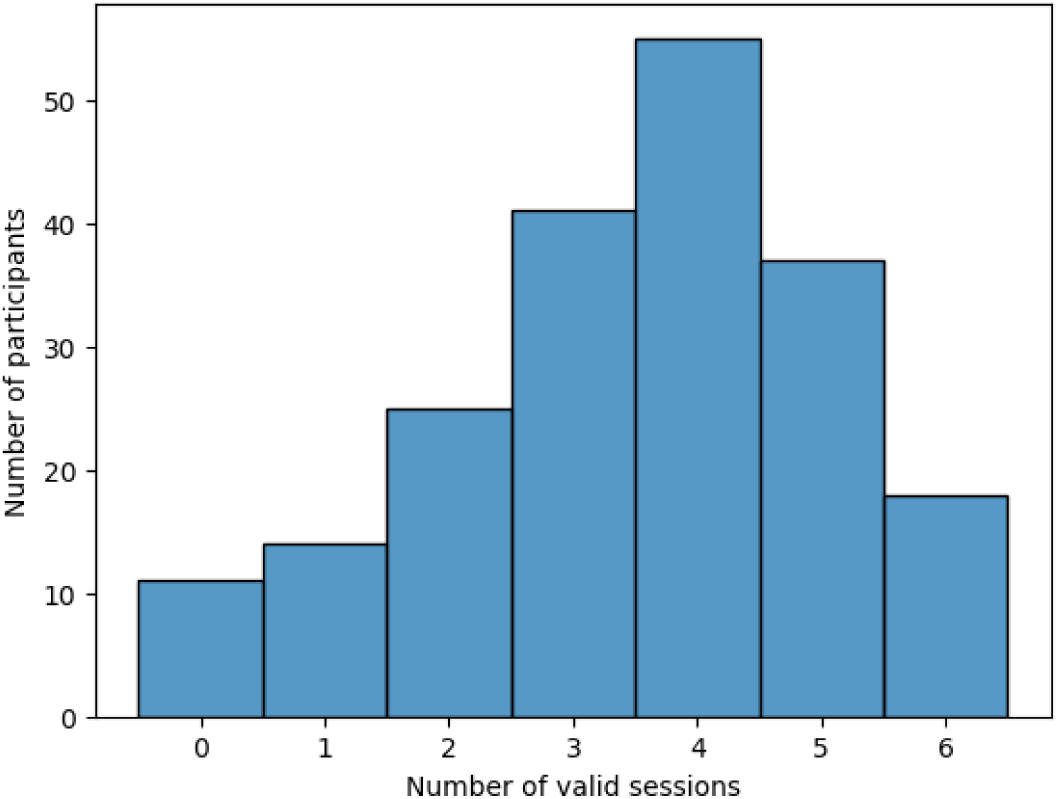
Histogram showing how many participants contributed valid data across the six NIRS sessions at 5, 8, 12, 18, 24 months and 3-5 years.

### 3.1 *First analysis*: profiling the fNIRS responses

To study the hemodynamic response function (HRF) peaks we first investigated the time-to-peak for the different conditions at different ages (Figure 5).

**Figure 5.**
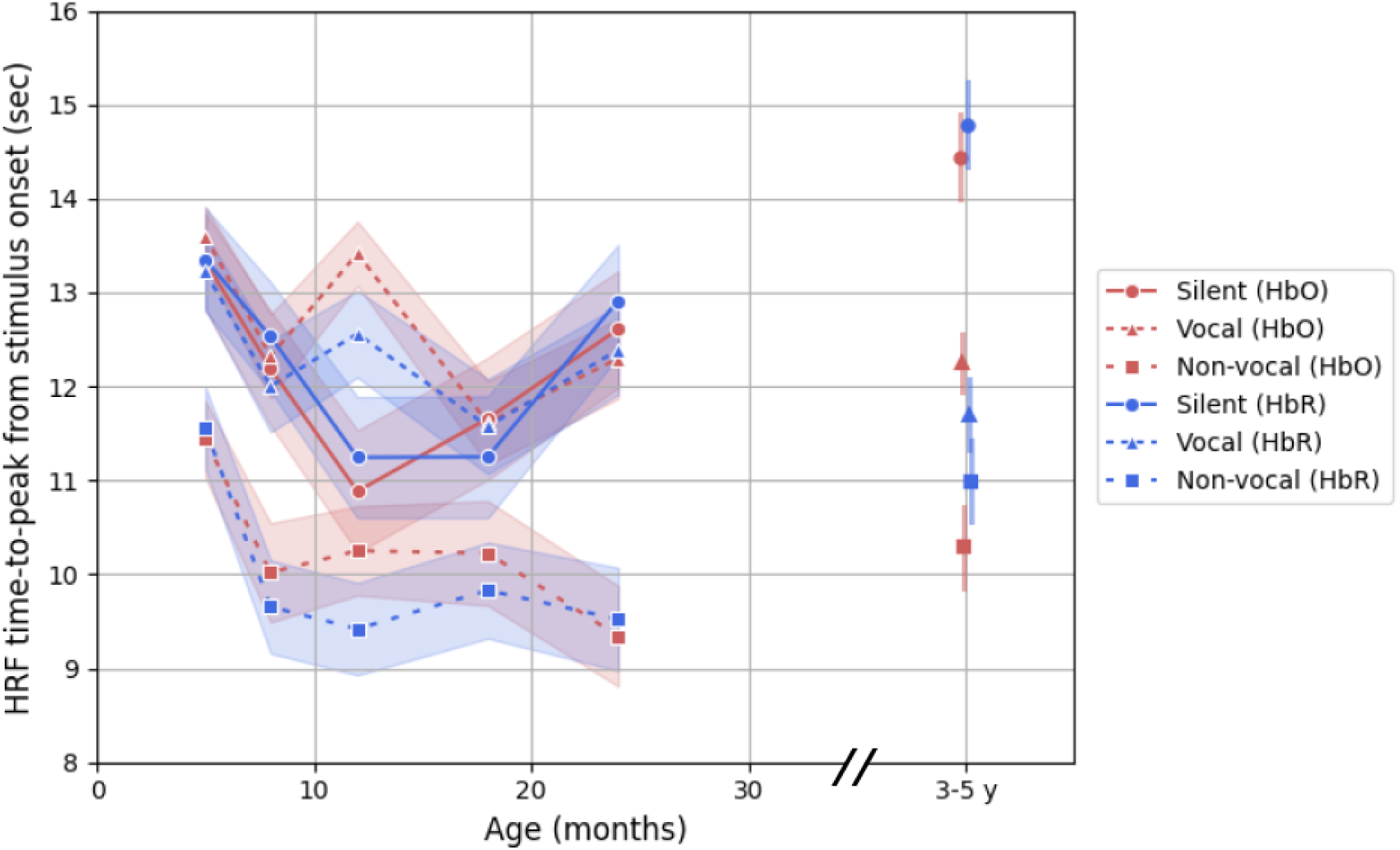
Time-to-peak from stimulus onset for the visual social (silent), auditory social (vocal) and auditory non-social (non-vocal) conditions (HbO and HbR) across ages (brands represent the standard error across participants).

The two-way ANOVA (for the condition and age point) for HbO and HbR showed significant effects of both the conditions and the age points (all p-values < 0.05). These results led us to use, for the study of activation, an approach centring the HRF window averaging around the mean time to peak (as opposed to a constant set time window). Therefore, a 4-second time window centred around the average HbO time-to-peak across channels and cohort for each condition and at each age point was selected to compute the average haemoglobin concentration changes for each trial for further statistical analysis. For the analysis of the contrast between *auditory social (vocal)* and *auditory non-social (non-vocal)*, the average time-to-peak across the two conditions was used to centre the time window used to extract the mean HRF concentration changes, which ranged from 8.8-12.8 seconds post-stimulus onset to 10.5-14.5 seconds depending on the age points (Table 2). For the analysis of the *visual social (silent)* condition, the average time-to-peak on that condition was used, which resulted in a time windows ranging from 8.9-12.9 seconds post-stimulus onset for the earliest to 12.4-16.4 seconds for the latest (Table 2).

**Table 2.** Time window around the average HbO time-to-peak used across different age points in seconds post-stimulus onset for the auditory social (vocal) > auditory non-social (non-vocal) contrast and visual social (silent) condition.

| Session | Auditory vocal > non-vocal | Visual social (silent) |
| --- | --- | --- |
| 5 mo | 10.5-14.5 seconds | 11.3-15.3 seconds |
| 8 mo | 9.2-13.2 seconds | 10.2-14.2 seconds |
| 12 mo | 9.8-13.8 seconds | 8.9-12.9 seconds |
| 18 mo | 8.9-12.9 seconds | 9.7-13.7 seconds |
| 24 mo | 8.8-12.8 seconds | 10.6-14.6 seconds |
| 3-5 years | 9.3-13.3 seconds | 12.4-16.4 seconds |

We also studied the absolute peak amplitudes of the HbO and HbR HRFs across age points for which a graphical representation can be seen in Figure 6. The two-way ANOVA tests for HbO and HbR did show a significant effect of both the conditions and the age points on the absolute maximum magnitude of the HRFs (all p-values smaller than 0.01).

**Figure 6.**
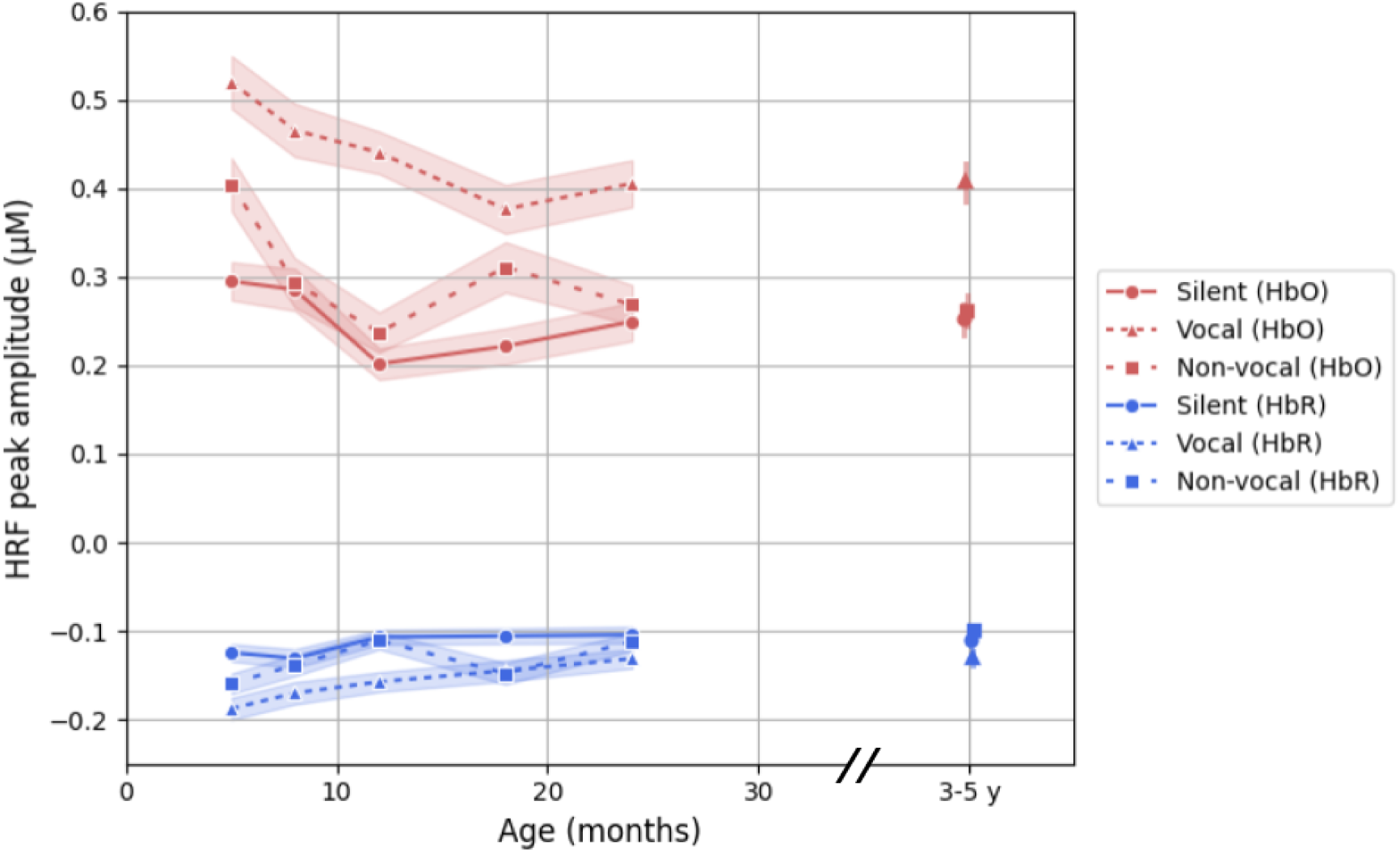
Value of the peak (in µM) for the HRF of the visual social (silent) auditory social (vocal) and auditory non-social (non-vocal) conditions (HbO and HbR) across ages (in months) compared to the pre-trial baseline of 2 seconds (bands represent the standard error across participants).

### 3.2 *Second analysis*: cross-sectional group level analysis

#### Visual social (silent)

With group-level analysis at each of the age points (channel by channel t-tests, FDR corrected), we found that the visual social stimuli elicited activation in channels covering regions known to be involved in social perception across all age points. The topographic map of brain activation can be seen in Figure 7. Overall, the number of significant channels decreased during infancy between *5* and *24 months*, suggesting the response becomes more localised over the posterior middle/superior temporal gyri and inferior frontal gyri. 13 channels showed statistically significant activations at *5 months*, 8 channels at *8 months* and 1-4 between *12-24 months*. Interestingly, the number of channels activated at *3-5 years* increased to 20 channels. Results showed a selective response for the *visual social (silent)* stimuli over the right posterior temporal region across all six time points from *5 months* to *3-5 years* of age (with 4 of 6 ages also showing activation in this region of the left hemisphere). Inferior frontal regions showed age-related changes in brain activation with the youngest (≤ *8 months*) and oldest (*24 months* to *3-5 years*) time points evidencing selective social visual responses. In fact, just under 50 % of the active channels at *3-5 years* were found in inferior frontal regions, which drove the pattern of developmental change to an increase in the overall number of channels active at this age.

**Figure 7.**
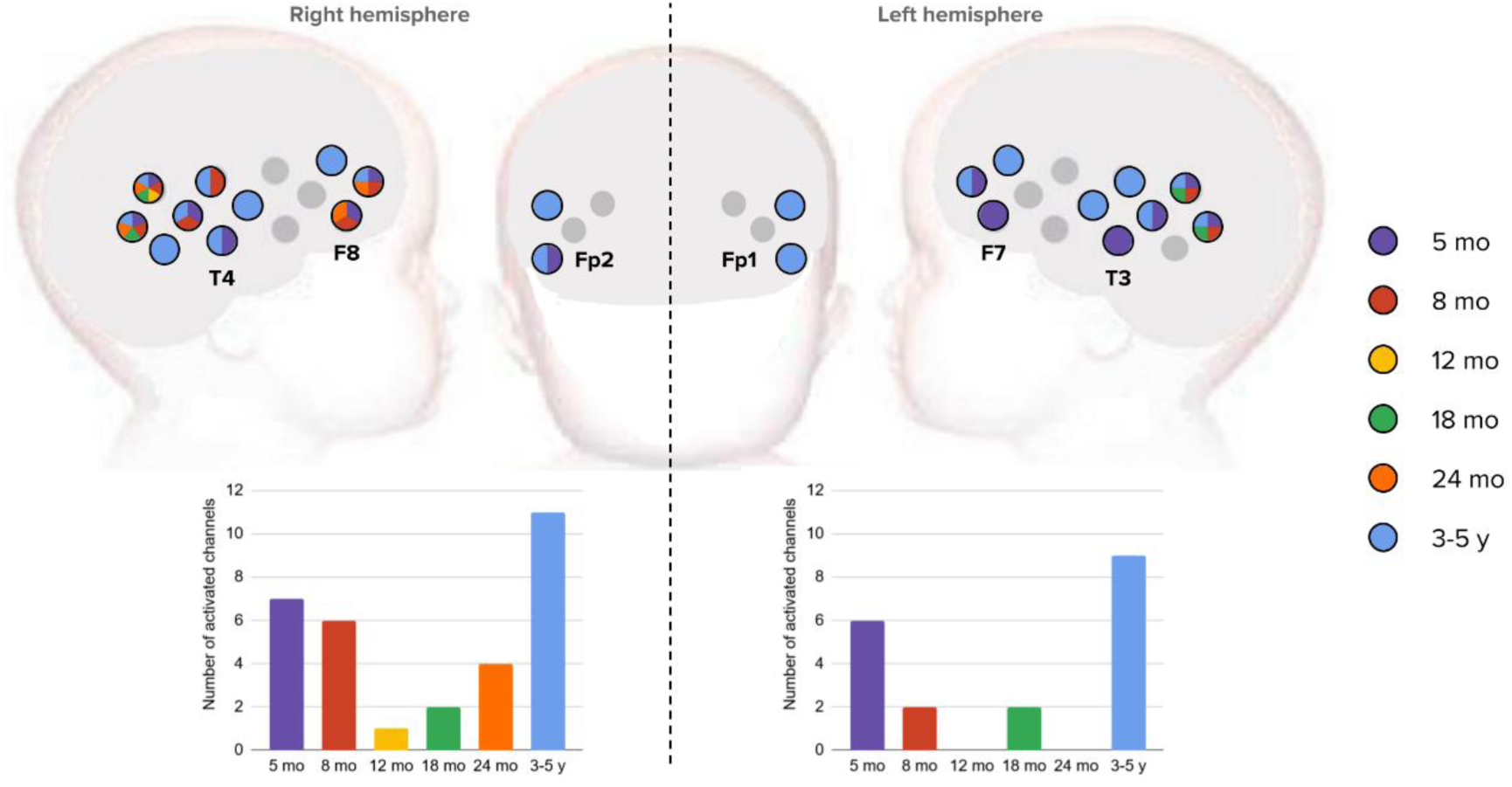
Graphical representation of the channels that showed significant activation to the visual social (silent) condition (significant increase in HbO and/or significant decrease in HbR). Each channel location shows the age points for which significant activation was detected (see legend for colour coded age-related responses). The bar charts represent the evolution of the number of channels with this significant contrast activation for each hemisphere across ages.

#### Auditory social (vocal) versus non-social (non-vocal)

For the contrast between *auditory social (vocal)* and *auditory non-social (non-vocal)*, stimuli (using paired t-tests, FDR corrected), we observed greater activation in response to vocal compared to non-vocal stimuli across all ages.

Contrary to the social visual selective responses, auditory selective responses revealed a stronger and more prolonged response to vocal compared to non-vocal stimuli across all six age points from *5 months* to *3-5 years* old.

Initially, at *5 months*, 8 channels showed a greater response to *auditory social (vocal)* than *auditory non-social (non-vocal)*, increasing to 10 at *8 months* and 27 at *12 months*. The number of channels with a significant contrast then decreased to 3 at *18 months* and increased again to 23 and 21 channels at *24 months* and *3-5 years* respectively.

The selective response was observed in the temporal regions, especially around the superior temporal gyrus bilaterally, with an area of selective activation growing up at *8 months* and *12 months*, where we also observed a response in the inferior frontal gyri bilaterally. The selective response became much more localised at *18 months*, and became again more widespread around the superior temporal gyrus at *3-5 years*. The topographic representation of activation can be seen in Figure 8. The bilateral anterior temporal regions show consistent activation across all six time points, localised in the anterior superior and middle temporal gyri. We also studied the inverted contrast which showed *no auditory non-social (non-vocal)* response greater than *auditory social (vocal)* for any of the channels at any of the ages from *5 months* to *3-5 years* (the graphical representation can be seen in Appendix 2). Examples of hemodynamic responses across ages can be found in Figure 9, with a channel in the right superior temporal gyrus that showed a significantly greater response to *auditory social (vocal)* than *auditory non-social (non-vocal)* at all sessions except *18 months* (Figure 9a). Note however that two neighbouring channels in this hemisphere did show social selectivity at *18 months*, and the homologous channel in the left hemisphere also revealed significant selectivity at that age. We also show in Figure 9b the hemodynamic responses of a left inferior frontal gyrus channel with a significant contrast only at *24 months* and *3-5 years*. While some portions of the time courses may look different across conditions, significant differences on the contrasts were considered only on the 4-second time windows specific to each age (see Table 2) to avoid bias towards any of the conditions.

**Figure 8.**
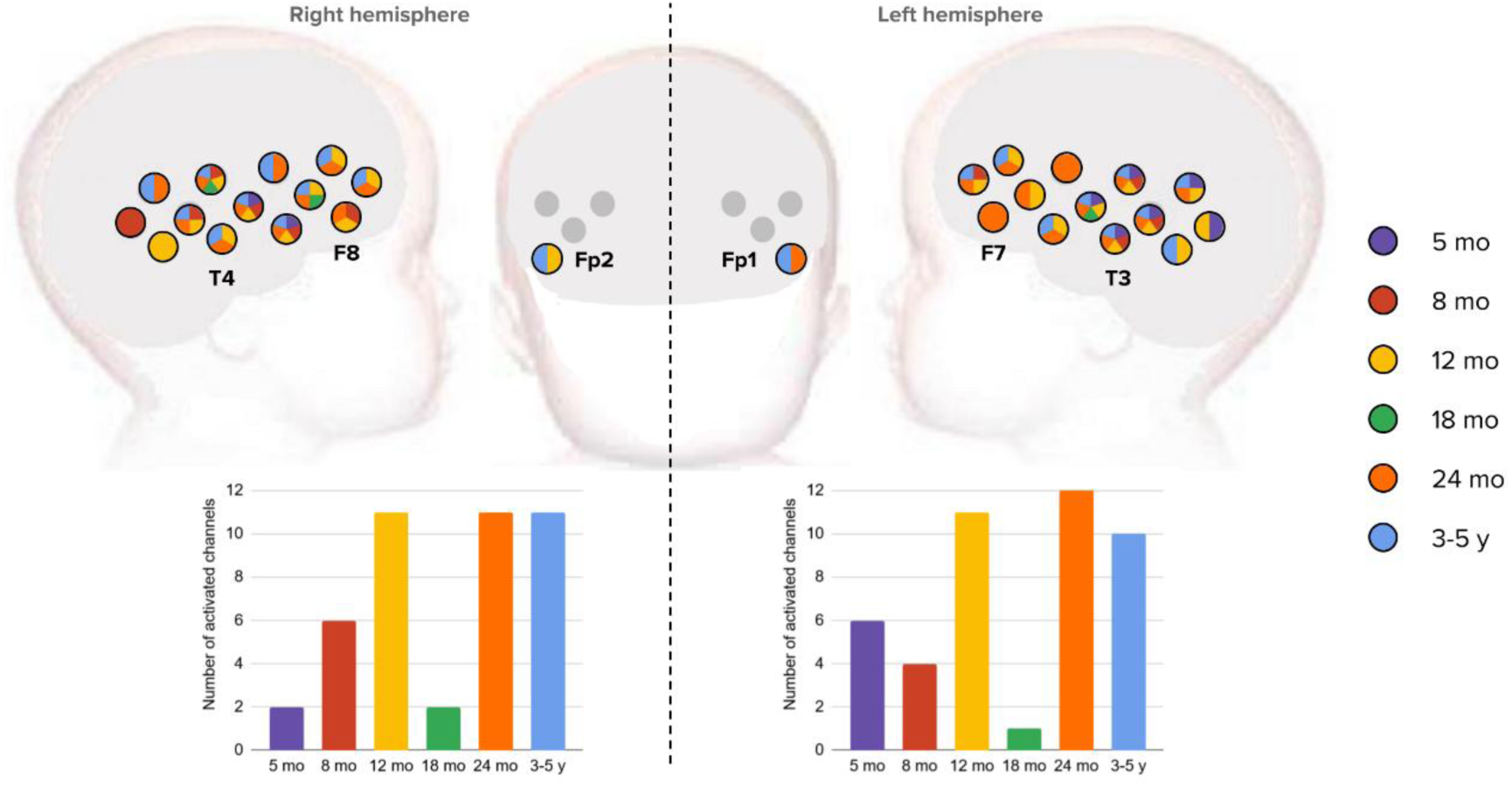
Graphical representation of the channels that showed significant activation on the auditory social (vocal) > auditory non-social (non-vocal) contrast (significant increase in HbO and/or significant decrease in HbR). Each channel location shows the age points for which significant activation was detected (see legend for colour coded age-related responses). A mask was applied such that we considered a significant activation on the contrast only if there was a significant activation to auditory social (vocal) compared to baseline in the first place. The bar charts represent the evolution of the number of channels with this significant contrast activation for each hemisphere across ages.

**Figure 9.**
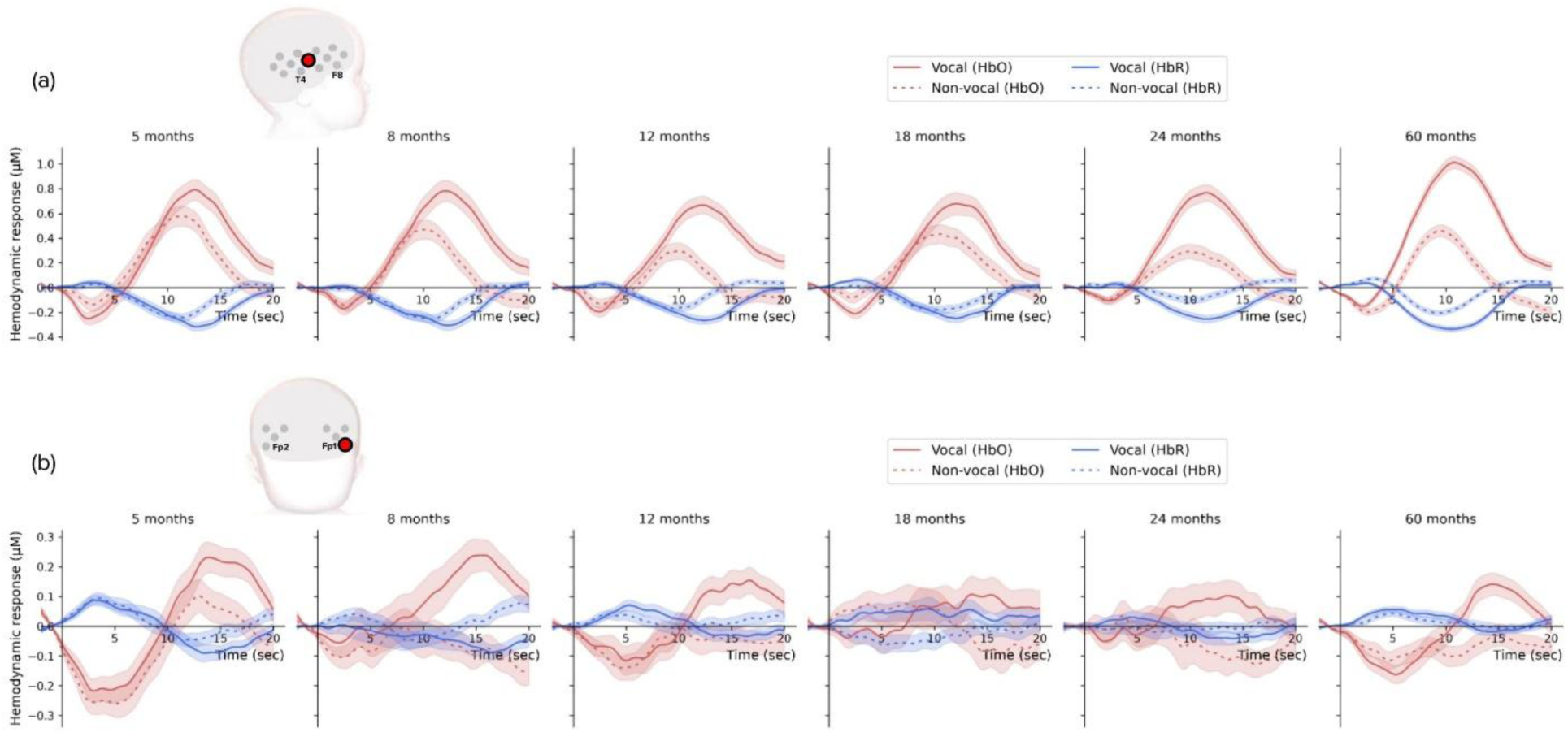
Examples of hemodynamic responses for the auditory social (vocal) and auditory non-social (non-vocal) conditions across ages, (a) in a right superior temporal gyrus channel which revealed social selectivity at all ages except 18 months, and (b) in a left inferior frontal gyrus channel which revealed social selectivity at 24 months and 3-5 years. Coloured shade bands represent the standard error.

### 3.3 *Third analysis*: longitudinal analysis of individual cortical specialisation

#### Individual social selectivity responses across ages

We visualised the proportion of participants showing social or non-social selectivity at each age point (Figure 10). This demonstrates that a greater proportion of participants showed selectivity towards social stimuli as they grew older: from 55% of the participants showing social selectivity at *5 months* to 77% at *3-5 years*. When having a closer look at the data to try to understand how participants moved between the different selectivity groups across ages, out of the 41 infants with an *auditory non-social (vocal)* selectivity at *5 months*, we saw that most of them (65.9 %) showed vocal selectivity in the majority of sessions after (from *8 months* to *3-5 years*). We also observed that only 2 participants showed only non-social selectivity or no selectivity at every age point from *5 months* to *3-5 years*.

**Figure 10.**
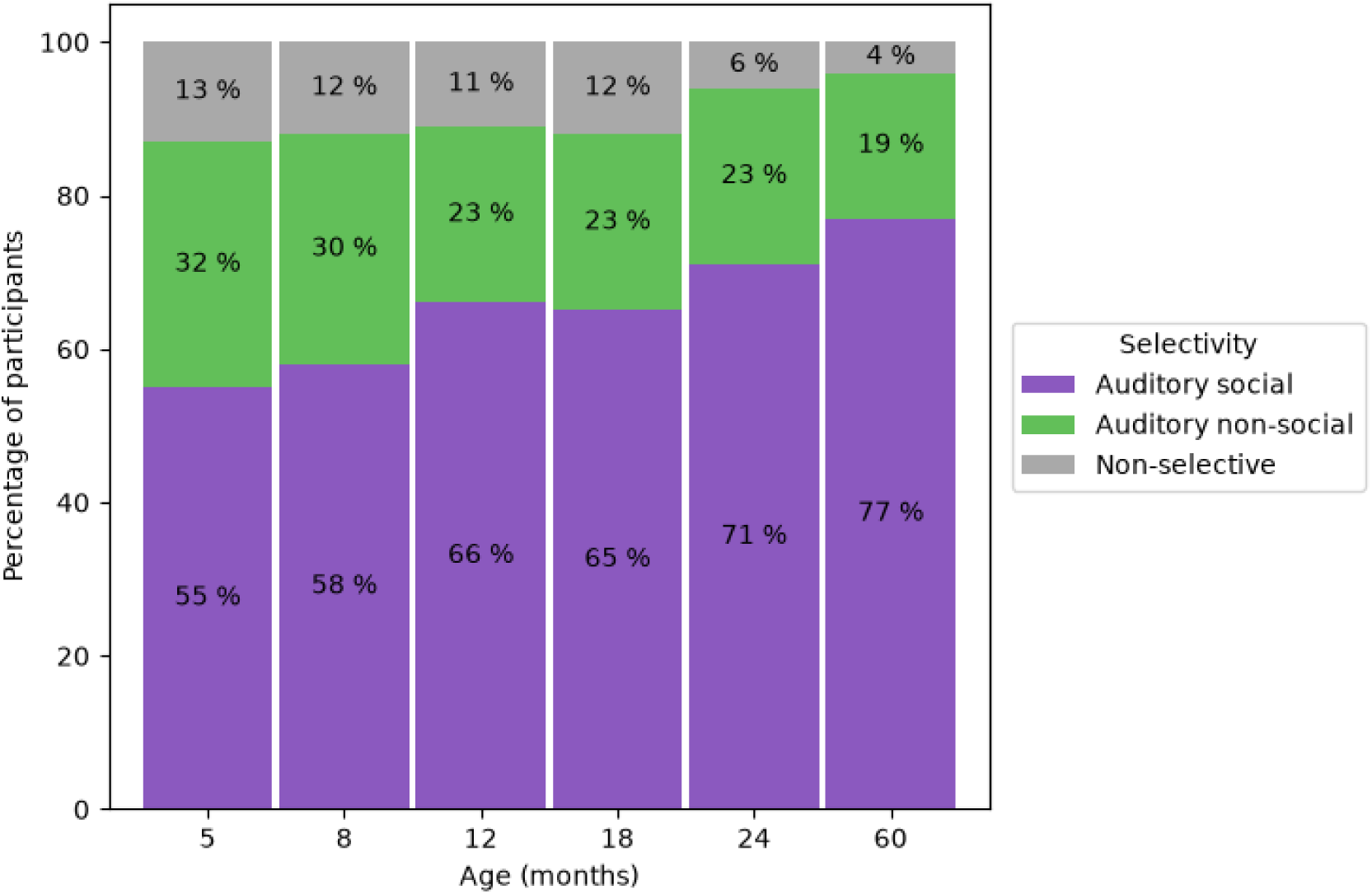
Bar plots showing the number of participants who showed selectivity towards each condition at each age point (auditory social (vocal) selectivity, auditory non-social (non-vocal) selectivity, or no selectivity). Selectivity towards a condition is defined as an HbO activation in response to that condition greater than the other condition in the anterior temporal regions of interest. Participants who did not have HbO activation in response to any of the conditions were represented as non-selective. Number of participants: 127 at 5 months, 110 at 8 months, 116 at 12 months, 111 at 18 months, 112 at 24 months, 124 at 3-5 years.

In studying the relationship of social selectivity with age with the Bayesian GLMM (Table 3), we found that the posterior mean for the fixed effect of age was 0.0246 with a posterior standard deviation of 0.0037 which suggests that an increase in age is associated with a higher probability of a social selectivity. The 95% credible interval for age (fixed effect) did not include zero, indicating that this effect is statistically significant. Additionally, the intercept term had a posterior mean of 0.2312 with a posterior standard deviation of 0.0831 which suggests that, at even before 5 months the log-odds of a social selectivity are slightly positive. Finally, the random intercepts for participants (random effect) showed a posterior mean of -0.4700 with a posterior standard deviation of 0.0513. The standard deviation of the random effects was estimated to be 0.625, with a 95% credible interval ranging from 0.564 to 0.693. This indicates substantial variability in the intercepts across different participants.

**Table 3.** Posterior estimates for the Bayesian Mixed Generalised Linear Model with a binomial link function, showing the relationship between age and social selectivity with participants as random effect. The fixed effect of age quantifies the change in log-odds per unit increase in age, and the random intercepts account for variability across participants.

| Parameter type | Posterior mean | Posterior standard deviation | Standard deviation | Standard deviation (lower bound) | Standard deviation (upper bound) |
| --- | --- | --- | --- | --- | --- |
| Intercept | 0.2312 | 0.0831 |  |  |  |
| Age (months) | 0.0246 | 0.0037 |  |  |  |
| Participant intercept | -0.4700 | 0.0513 | 0.625 | 0.564 | 0.693 |

We then investigated the 50 participants with data at all sessions during the first year of life with Figure 11 showing the HbO responses for the *auditory social (vocal)* and *auditory non-social (non-vocal)*, which enabled us to compare the response to the conditions for each participant specifically (link between data points).

**Figure 11.**
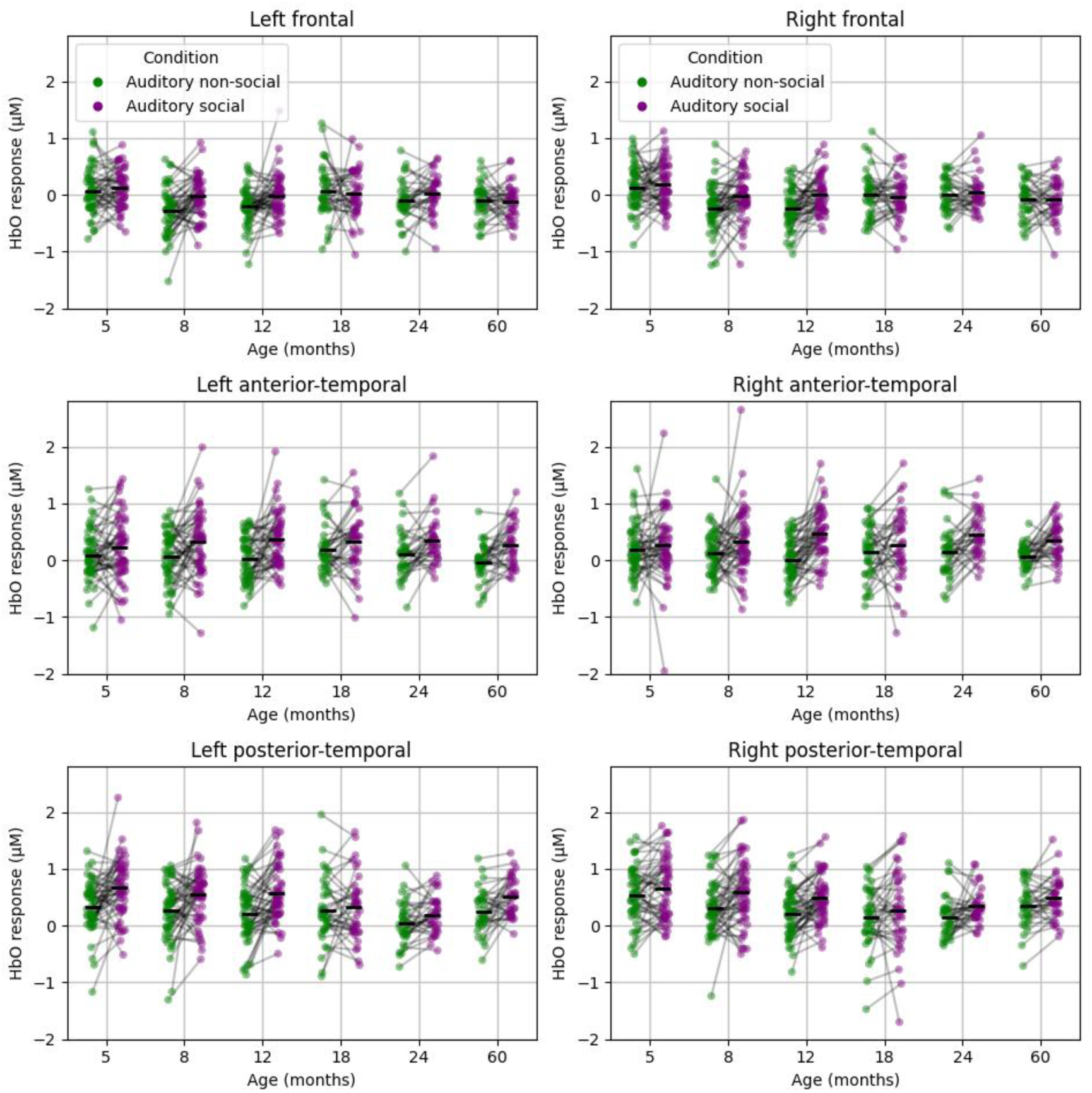
Graphical representation of the auditory social (vocal) and auditory non-social (non-vocal) HbO responses in the different regions of interest for each age point. Green dots represent individual auditory non-social (non-vocal) HbO responses while purple dots represent auditory social (vocal) HbO responses, with black marks representing the group averages. Links between the data points represent individuals.

The most consistent patterns of auditory social selectivity across participants were observed in anterior temporal regions (middle panels of Figure 11), as most participants showed an increase between *auditory social (vocal)* and *auditory non-social (non-vocal)*. This was different than in the posterior temporal regions (bottom panels of Figure 11) where the participants showed more varied trends.

More precisely, in the left anterior temporal region of interest, 60 % of participants showed a stronger response to *auditory social (vocal)* than *auditory non-social (non-vocal)* at *5 month* (HbO average response of 0.23 µM for *auditory social (vocal)* and 0.09 for *auditory non-social (non-vocal)*), 66 % at *8 months* (averages respectively 0.32 and 0.07 µM), 74 % at *12 months* (0.37 and 0.02 µM), 65 % at *18 months* (0.34 and 0.19 µM), 76 % at *24 months* (0.36 and 0.10 µM), 68 % at *3-5 years* (0.28 and -0.03 µM).

In the right anterior temporal region of interest, 54 % of participants showed a stronger response to *auditory social (vocal)* than *auditory non-social (non-vocal)* at *5 month* (HbO average response of 0.26 µM for *auditory social (vocal)* and 0.19 for *auditory non-social (non-vocal)*), 60 % at *8 months* (respectively 0.34 and 0.13 µM), 84 % at *12 months* (0.47 and 0.00 µM), 57 % at *18 months* (0.27 and 0.14 µM), 76 % at *24 months* (0.45 and 0.14 µM), 71 % at *3-5 years* (0.35 and 0.06 µM).

#### Developmental trajectories

Finally, to further explore individual patterns of *auditory social (vocal)* > *auditory non-social (non-vocal)* we investigated the developmental trajectories of the 50 participants who had data at all sessions from *5* to *12 months* (Figure 12), with participants divided into groups based on the first age point of social selectivity (*auditory social (vocal)* > *auditory non-social (non-vocal)*) in the temporal anterior regions of interest. Of those 50 participants, 26 had their first social selectivity at *5 months* (52.0 %), 13 at *8 months* (26.0 %), 8 at *12 months* (16.0 %), and 3 with no social selectivity by *12 months* (6.0 %). The group of participants with first social selectivity at 5 months showed very clear responses to social stimuli greater than non-social stimuli at all the sessions. On the other hand, for participants with a first social selectivity at 8 and 12 months, we saw that even though they tended to remain overall socially selective after their first selectivity, the *18 months* session seemed to be an exception, as seen in the previous group results with the group level showing significant contrasts in fewer channels (Figure 8). Indeed, here there appeared to be a trend towards the inversion of the social selectivity at that age (non-social stimuli eliciting a greater anterior temporal brain response than social stimuli). Finally, the 3 participants out of the 50 with data at all sessions over the first year of life which did not show a social selectivity by age 12 months exhibited trajectories with much more variability.

**Figure 12.**
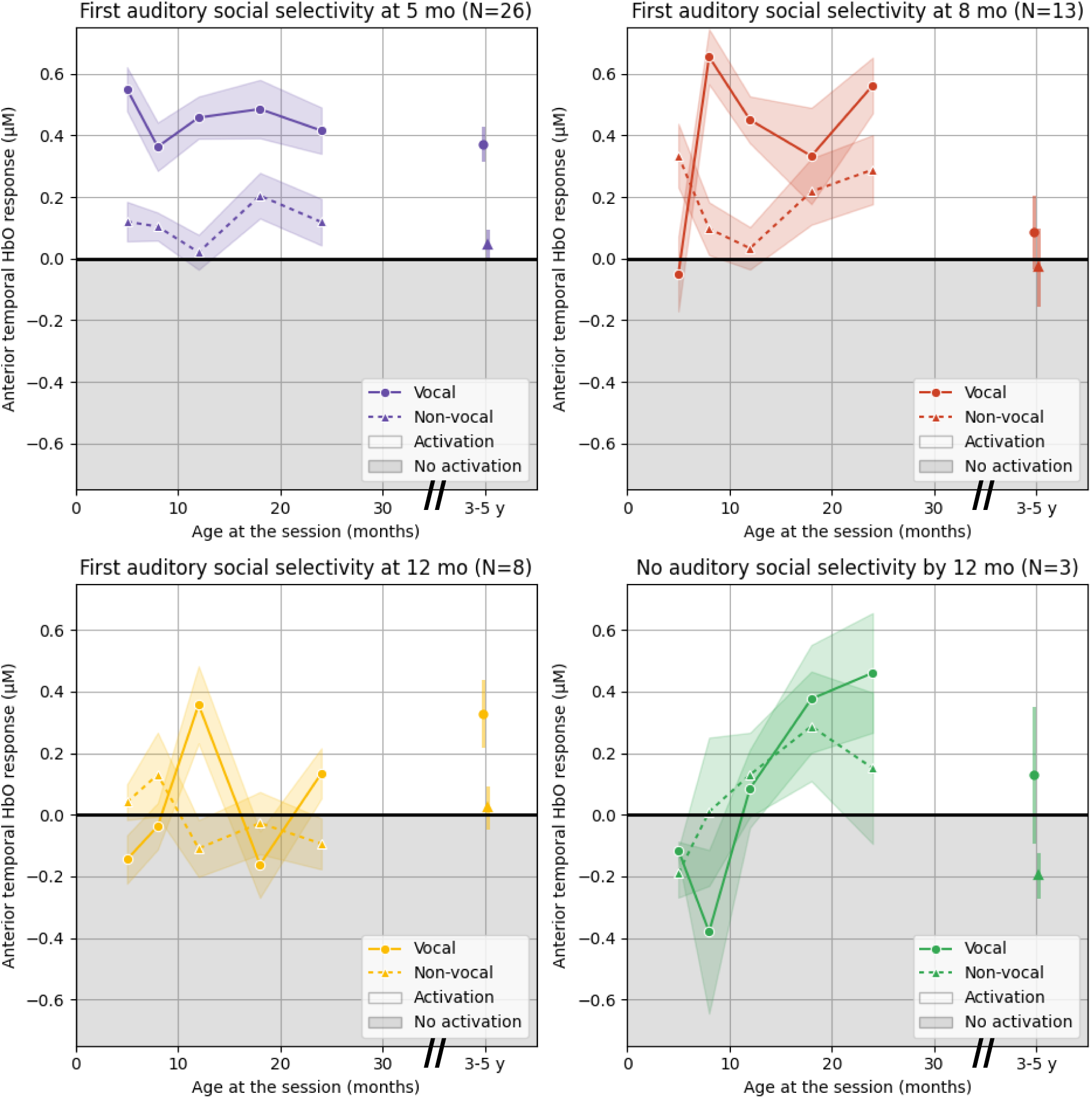
Graphical representations of the mean HbO responses (mean over a 4-second window centred around the cohort average peak response delays as described in Table 2) in anterior temporal regions for the auditory social (vocal) and auditory non-social (non-vocal) conditions across ages, for participants with data at all sessions between 5 and 12 months (N=50). Each panel represents a separate social specialisation profile group based on the age of first auditory social selectivity. Coloured shade bands represent the standard error.

## 4. Discussion

### 4.1 Social perception

This study, using data from a longitudinal study in rural Gambian infants, has shown that for visual social selectivity: ***(i)*** infants showed brain responses to visual social stimuli in the posterior temporal regions at *5*, *8*, *12*, *18*, *24 months* and *3-5 years* of age; ***(ii)*** a region of the right posterior temporal social brain network was consistently activated across all timepoints; ***(iii)*** additional inferior frontal brain regions were also activated but unlike temporal responses, different developmental patterns of activation were observed, with greater recruitment of this region when infants were younger (5-8 months), a drop in activation in mid-infancy, then a rise in late toddlerhood to pre-school age (2-5 years); and finally ***(iv)*** overall, when considering the period of infancy specifically (0-2 years), the responses changed from a more diffuse activation distributed across multiple channels within each inferior frontal and posterior temporal region in the younger time points to more localised activation within a narrower number of channels during the second year of life suggestive of developmental specialisation.

For auditory social selectivity (vocal > non-vocal): ***(i)*** responses were present across frontal and temporal regions at *5*, *8*, *12*, *18*, *24 months* and *3-5 years* and ***(ii)*** contrary to the visual social selective response, which revealed a narrowing of active channels across 0-2 years, auditory social selectivity at *5* and *8 months* was found in a smaller number of channels, which became more widespread across inferior frontal, anterior temporal and posterior temporal regions at 1-5 years. Furthermore, ***(iii)*** while the *18 months* age point was in line with the observed pattern of developmental specialisation in the visual social contrast, the auditory social selective response patterns were markedly different to the overall trajectory seen. Finally, the individual longitudinal infant analyses of the sub-sample (*N*=50) with valid data at all timepoints over the first year enabled us to group infants based on their age of selectivity towards auditory social stimuli in anterior temporal regions: selectivity emerging at *5 months*, *8 months*, *12 months* and later than *12 months*.

The visual social selectivity confirms previous studies from the UK, the Netherlands, The Gambia and Bangladesh also showing brain activation in the posterior temporal regions in infants from 4 to 6 months of age (Lloyd-Fox et al., 2009; 2011; 2013; 2014; 2017; 2018; Siddiqui et al., 2023; Braukmann et al., 2018; Perdue et al., 2019; Pirazzoli et al., 2022; Blanco et al., 2023; Frijia et al., 2021). Furthermore, in line with studies with adults (i.e. Pelphrey et al., 2005) several of these previous studies also find social visual selective patterns of inferior frontal brain activation to similar social stimuli (Blanco et al., 2023; Frijia et al., 2021; Lloyd-Fox et al., 2009; 2011; Perdue et al., 2019; Pirazzoli et al., 2022) confirming that this region of the brain is recruited at this stage of infant development. Evidence of visual social selectivity in the second year of life (Lloyd-Fox et al., 2017; Pirazzoli et al., 2022) and during pre-school years (Perdue et al., 2019; Pirazzoli et al., 2022) is less common. Interestingly, while many of the previous publications in young infancy are from cross-sectional studies conducted in European specialist research labs, previous research that extends longitudinally into the pre-school years (2- to 5-year-olds) has to date only been conducted by the current study and the study from Bangladesh. The current study builds on this evidence by integrating a significantly higher number and density of longitudinal time points into the protocol to better understand the developmental profile across the first five years of life. Firstly, this developmental trajectory of brain specialisation to visual social selectivity revealed greater recruitment of inferior frontal brain regions when infants are young (5-8 months) and then returning to this pattern again in late toddlerhood to pre-school age (2-5 years). This later reemergence of frontal responses is in line with previous findings from the Bangladesh cohort (Perdue et al., 2019; Pirazzoli et al., 2022). One possibility that could explain this additional recruitment of frontal regions in pre-schoolers is if the children are viewing the stimuli with more in-depth processing. It is plausible that they could be trying to understand and interpret the intentions or meaning of the actors in the videos in a more complex manner when they reach this age, thus recruiting higher attentional resources in frontal regions. This would align with previous functional magnetic resonance imaging (fMRI) work in adults showing recruitment of frontal cortical regions when interpreting more complex social abilities such as inferring others’ mental states (i.e., Saxe & Wexler, 2005) and interpreting intention in actions (i.e., Saxe et al., 2004). Currently further work is required to better understand social brain network function in the pre-school years as due to the number of challenges of undertaking neuroimaging studies in this age range, this remains an age period that is relatively under-investigated. The additional age points used in our study allowed a deeper understanding of the response profile across the first two years. In line with previous fMRI research in adults (Allison et al., 2000; Pelphrey et al., 2005), the posterior temporal area revealed a consistent socially selective response across all six time points. Furthermore, findings revealed that the visual social brain responses specialised from a broad region of temporal and frontal activation at *5-8 months* to more restricted clusters within the posterior temporal region by *12-18 months*. These findings replicate a previous study (Lloyd-Fox et al., 2017) from different cohorts measured in The Gambia with the same paradigm (though note that only the temporal cortex was measured in this work). One possibility is that regions of the social brain critical for detecting communicative situations specialise rapidly across the first two years, reflecting neural interactive specialisation (Johnson, 2001). As the individual ages, the brain response becomes more specialised (and less diffuse) within each cortical region as the regions orchestrate a network of connected responses (Johnson et al., 2009). This could help us to understand why 12-18-month-olds did not respond to these stimuli with an activation as widespread as the younger infants.

This study also showed auditory social selective (vocal > non-vocal) brain activation, with particularly consistent activation across all age points in the anterior temporal regions. This concurs with previous findings in fNIRS and fMRI studies at 4 to 7 months (Blasi et al., 2011; Grossmann et al., 2010; Lloyd-Fox et al., 2012, 2013; 2017; Minagawa-Kawai et al., 2011; Perdue et al., 2019; Pirazzoli et al., 2022; Shultz et al., 2014; Collins-Jones et al., 2024), the second year of life (Lloyd-Fox et al., 2017; Pirazzoli et al., 2022), pre-school 3-5 years (Perdue et al., 2019; Pirazzoli et al., 2022) and into adulthood in fMRI work (Belin et al., 2000; Pernet et al., 2015). While auditory social selectivity at *5* and *8 months* replicated previous findings of a localised anterior temporal response, the response became more widespread across inferior frontal, anterior temporal and posterior temporal regions at 1-5 years. In the fMRI literature with adults, wide patterns of voice specific activation are also found in the superior temporal cortex (Belin et al., 2000; Pernet et al., 2015; Frühholz et al., 2020). Another previous work describes widespread activation across these brain regions with fNIRS in a previous cohort in Bangladesh. This was observed consistently across two time points at 6 months and 2 years of age, though a different cohort measured at 3 years revealed a very different pattern of responses, with very weak and isolated to temporal regions (Perdue et al., 2019; Pirazzoli et al., 2022). It is possible, given the findings from our work in combination with the published literature, that this indicates that contextual factors can significantly impact individual variance in the developmental profile of this response. It has been noted in previous work that the 4-7 month auditory socially selective responses have considerably varied across studies, sometimes revealing left lateralised responses (Blanco et al., 2023; Lloyd-Fox et al., 2011; Minagawa-Kawai et al., 2011; Shultz et al., 2014), sometimes right lateralised (Lloyd-Fox et al., 2013), sometimes bilateral (Grossmann et al., 2010; Perdue et al., 2019). Furthermore, age related changes even within this small developmental window have been observed, explaining some of the individual variance (Grossmann et al., 2010; Lloyd-Fox et al., 2011). While we are lacking previous evidence for the developmental window from 1-5 years, there are two possible explanations for the differences observed across the current study and previous work. Firstly, it could be driven by methodological factors, which could have impacted the strength of the observed signal as well as the power to observe statistically significant responses. The sample sizes used in the current study from The Gambia and the previously published work from Bangladesh (Perdue et al., 2019; Pirazzoli et al., 2022) ranged from 111 to 155 participants for each age point at which the cohorts were analysed. Conversely, for the previous cross-sectional studies from The Gambia, UK, Germany, USA. and Japan (Grossmann et al., 2010; Lloyd-Fox et al., 2012, 2013; 2017; Minagawa-Kawai et al., 2011; Shultz et al., 2014) the sample sizes were much lower ranging from 12 to 33 participants. Therefore, the findings for those studies in the Gambia and Bangladesh with the higher samples, providing more statistical power, are likely more robust. While different stimulus contrasts (for example vocal emotional and non-emotional cues, speech, human, animal), processing methods and thresholding of statistical effects could have driven differences in the localisation of the regionally reported responses in some of this previous work, we can be confident that the stimulus presentation, pre-processing steps, statistical methods employed and statistical thresholds set across several of our own publications were very similar and could not explain the differences in activation locations, spread of response or condition differences (Lloyd-Fox et al., 2012; 2017, current study).

Therefore, while methodological considerations could have influenced some of these findings, the age-related differences in the spread of activation into regions beyond the widely reported anterior temporal response are more likely to be explained by contextual factors that could impact an individual’s response. For example, why extensive activation in frontal-temporal regions were seen in the current study from 5 month to 5 years and in one Bangladesh cohort at 6 months and 2 years (whilst the second Bangladesh cohort measured at three years reports a very isolated small temporal response). Of interest, we observed a sharp decrease in the spread of activation at *18 months*, with only 3 bilateral temporal channels showing activation on the contrast versus 27 at *12 months* and 23 at *24 months*, which we cannot attribute to lower data quality at this age. Indeed, the anterior temporal location of the auditory socially selective response at *18 months* aligns with the most consistent response at all other ages as well as with previous work. This is very similar to the Bangladesh study where only 3 isolated bilateral channels showed activation at 3 years (Pirazzoli et al., 2022), also within these regions. While the response in the Bangladesh work comes from a distinct cohort and may represent individual variance in group level responses, it is more difficult to explain in our longitudinal study. Additional studies would need to be conducted in order to investigate the unexpected auditory social selectivity findings at *18 months*.

It is possible that insights gained from the individual participant neural profiles will help us understand the level of individual variance. Previous work has shown that 0-9 months is a period of transition, where some infants may respond more strongly to non-vocal stimuli while others to vocal stimuli (i.e. Blanco et al., 2023; Lloyd-Fox et al., 2012). In the current study, we identified the age at which infants started showing a greater anterior temporal activation in response to *auditory social (vocal)* compared to *auditory non-social (non-vocal)* stimuli. The proportion of infants already vocal selective at *5 months* on the restricted sample with all sessions from *5* to *12 months* (50 participants) was 52 %, which is close to what we see on the larger sample with the 127 participants with valid cross-sectional data at the *5 months* session (55 %). Furthermore, two thirds of these infants continued to show consistent vocal selectivity at later age points. This, however, did not mean that their selectivity was already significant, as we were not able to test this statistically at the individual level due to the limited number of trials per condition for each participant (as low as 3 for some infants). Nonetheless, we believe it was still useful to build such group-based thresholds of participants’ block-averaged contrast responses, as it enabled us to have an indication of their specialisation trend. The age of selectivity in our study did appear to be in accordance with previous literature, highlighting variance in vocal selectivity at 4 to 8 months (Grossmann et al., 2010; Lloyd-Fox et al., 2012), which becomes more consistent at later ages. It seemed established, however, that while a temporal auditory response is detected during the first week of life the development of selective social perception at this age is not yet mature, as shown by Cristia et al. (Cristia et al., 2014), and reveals considerable individual variation (see Greenhalgh et al., 2025 for analyses at 1 months from the BRIGHT project). While the significant positive effect of age on the probability of auditory social selectivity suggests in our study that older participants are more likely to show social selectivity, the substantial variability in the random intercepts indicates that there are individual differences that are not captured by the fixed effects alone. This highlights the importance of building different profiles of social specialisation, which supports our proposal of an approach to group participants based on the age of first social selectivity, showing different developmental trajectories for brain responses to social and non-social stimuli. Those different social specialisation profiles (subgroups) will become useful for future work to see if they could be linked to adversity or resilience factors. In the Bangladesh study, differences in activation patterns for both the visual social selective responses and the auditory selective responses were partially explained by age dependent associations with contextual factors. Cortical social discrimination at 2 years related to family reported intimate partner violence, verbal abuse and family conflict, while at 3 years it related to verbal abuse and family conflict and maternal depression (Pirazzoli et al., 2022). No associations were found with the younger age point of 6 months. In future work we aim to model the current findings with broader contextual factors prevalent in the rural population where the BRIGHT Project has taken place (Lloyd-Fox et al., 2024) to explore the impact of both poverty associate risk and enriched family contexts.

### 4.2 Methodological consideration

Studying HRF characteristics was very important here, as we saw that both the time-to-peak and magnitude of the HRFs were affected by the experimental condition and age. It enabled us to use adaptive delays for the time windows used for signal averaging, which prevented introducing bias towards an age, a participant, a brain region, or a condition.

The investigations of those HRF characteristics led us to conclude first of all that because the time-to-peak was influenced by the stimuli condition (Figure 5), it was important to select a time window for feature extraction that was centred around the average time-to-peak across the compared conditions. Secondly the fact that the time-to-peak was influenced by the age of participant (Figure 5) added confidence that the time-to-peak selected for window centring should also be age dependent and therefore calculated separately for every age point. Finally, the fact that age was also influencing the magnitudes of the responses (Figure 6), combined with the growing head circumference throughout infancy (Table 1) led us to forgo using HRF amplitude features directly for any comparisons across age points. Instead, we focused on the sign of the contrast (selectivity), rather than comparing values directly. While the differences in maximum amplitude cannot all be attributed to neural correlates alone and could originate for example from differences in skull thickness (Li et al., 2015), it remains nonetheless very important to keep this in mind, especially when comparing results at different age points. This is why for the longitudinal analysis, the focus was on relative changes between conditions (social or non-social selectivity) and age of first social selectivity, rather than HbO or HbR amplitude values.

Selecting a time-window for extracting and HRF average amplitude is a very common practice in fNIRS literature. This is especially the case for studies with infants because other approaches such as general linear model (GLM) are harder to use with infant populations (Filippetti et al., 2023). While a fixed window selection could work when doing cross-sectional studies (Gemignani et al., 2021), we showed here that it was very important to consider adaptive latencies when stimuli are different and the study is longitudinal.

### 4.3 Recommendations for individual differences analyses for future work

Based on our work, we propose the following recommendations:

1. We recommend exploring the anatomical spread of the response across the network of regions and not just the localisation of the peak response as this can reveal important age- or stimulus-related differences.
2. We also recommend exploring the latency of the response to different stimuli and at different ages to ensure the use of an analytical approach that can accommodate age- and stimulus-related changes that may be theoretically important.
3. Finally, future research should focus on using multiple longitudinal time points to understand not only if the responses are appropriate for a given age point, but also if deviations from a trajectory at the individual level are transient or related to a change in cognitive processing that is theoretically important.

## 5. Conclusions

This study contributes to the understanding of the development of social perception in infancy by examining brain responses to *auditory social (vocal)*, *auditory non-social (non-vocal)* and *visual social (silent)* stimuli longitudinally from *5 months* to *3-5 years* of age. Our findings indicate infants have a specialised neural response towards social stimuli, even as early as 5 months of age. Specifically, we observed a consistent posterior temporal selectivity for social visual stimuli across all ages, with some changes in inferior frontal regions, showing selectivity in the youngest and oldest age groups. Social vocal stimuli elicited a widespread and sustained response across temporal areas in this longitudinal cohort, extending to inferior frontal areas from 12 months. Moreover, our longitudinal analysis revealed distinct developmental trajectories in individual infants’ specialisation to social vocal stimuli, with infants generally remaining social selective after they showed their first social specialisation.

These results highlight the complexity of the developmental changes in the neural processing of auditory social information during infancy and early childhood and open to future research to investigate how these trajectories may be influenced by environmental factors (adversity or resilience factors) and how they relate to later social and cognitive outcomes. The methodology developed in this study, which combines the analysis of the latency and magnitude of the hemodynamic response with the characterisation of individual developmental profiles, can be valuable for future studies in developmental cognitive neuroscience, especially in under-represented populations and in low-resource settings.

## 6. Data and code availability

### 6.1 Data

As described in detail in the BRIGHT Project publication describing the full study protocol (Lloyd-Fox et al., 2024), access to any data collected during or generated by the BRIGHT Project is fully audited, and to ensure data security, is overseen by the data management team in the UK and The Gambia. While data sharing is critically important to maximising the benefit of research, the BRIGHT Project must also consider the need to protect the confidentiality of this sensitive group (particularly the infants within the mother-infant dyads, who as minors do not consent for themselves). The dataset is unique in that it provides the ability to link data points together (i.e. fNIRS data with outcome data or contextual factor data), but due to the nature of the data being collected (i.e. collected from a specific geographical location, longitudinal dataset of several datapoints) the majority of the data cannot be fully de-identified under the guidance included in the European General Data Protection Regulation (GDPR), and the conditions of our ethics approval do not allow public archiving of pseudonymised study data. Indeed, while fNIRS recordings can be anonymised, some of the demographic information used in this work cannot be deidentified.

However, collaborations are encouraged, and projects are evaluated primarily on their consistency with the ethical principles and aims of the project that the families signed up to when partaking in this study. To access the data, interested readers should contact the BRIGHT coordinator on our website (https://www.globalfnirs.org/contact). Access will be granted to named individuals following ethical procedures governing the reuse of sensitive data. Requestors must complete and sign a data sharing agreement to ensure data is stored securely and used in accordance with the BRIGHT Project’s policies on Ethics, Data Sharing, Authorship and Publication.

### 6.2 Code

The data analysis code is available on GitHub (https://github.com/globalfnirs).

## 7. Author contributions

JB: conceptualisation, data analysis and manuscript writing; CB: conceptualisation, data analysis and manuscript editing; EM: data collection; BB: data analysis and manuscript editing; ET: data collection; SMC: data collection; BM: project supervision and manuscript editing; SEM: project lead and manuscript editing; CE: project lead and manuscript editing; AB: conceptualisation, data analysis, manuscript editing and project supervision; SLF: conceptualisation, project lead and supervision, data analysis and manuscript writing.

## 10. Acknowledgements

We would like to thank the BRIGHT Project Team for their continued support and commitment to the research programme that enabled this publication. In particular, Giulia Ghillia and Marta Perapoch Amado for their help in project administration and behavioural coding. We also thank the families who took part in this study.

## Appendices

**Appendix 1.**
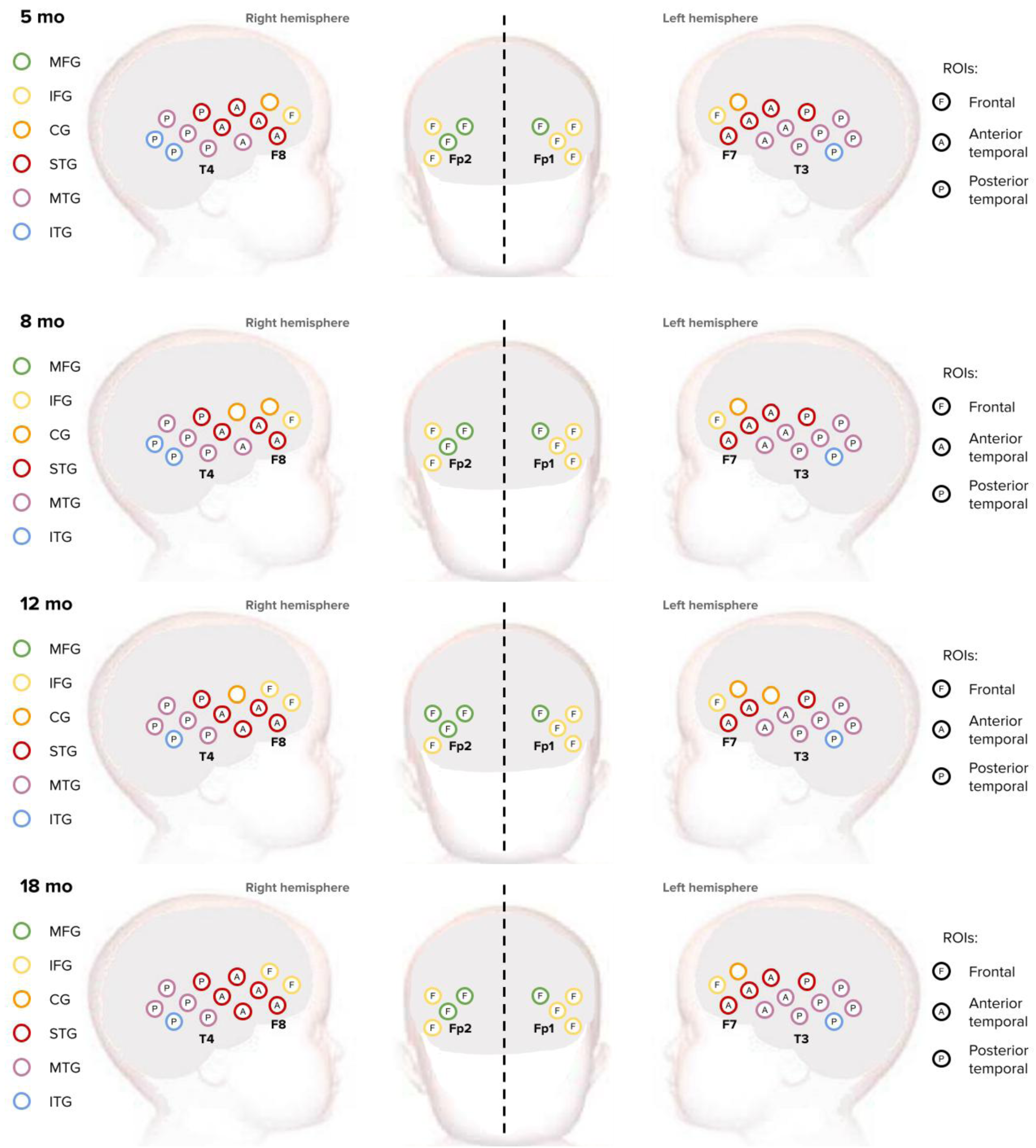
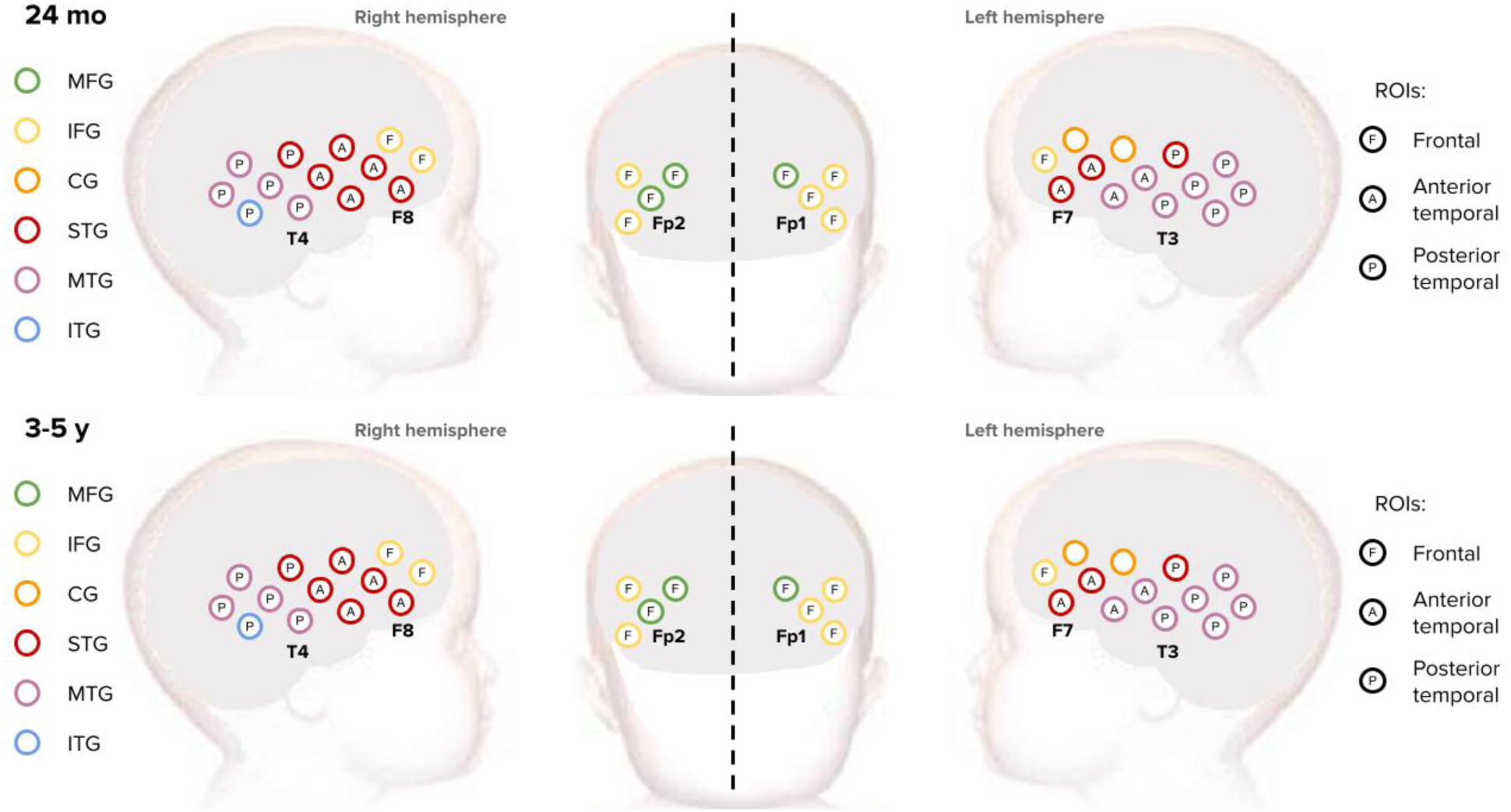
Regions of interests used for the longitudinal analysis (frontal, anterior temporal, posterior temporal) along with the anatomical co-registration of optodes: middle frontal gyrus (MFG), inferior frontal gyrus (IFG), pre/postcentral gyrus (CG), superior temporal gyrus (STG), middle temporal gyrus (MTG), inferior temporal gyrus (ITG).

**Appendix 2.**
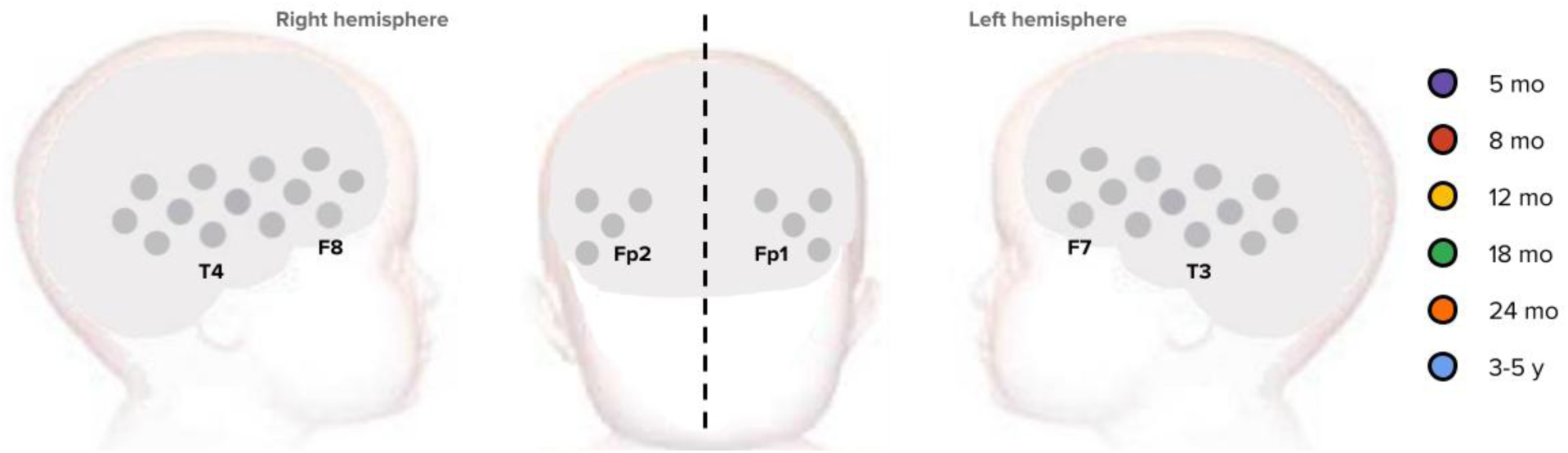
Graphical representation of the channels that showed significant activation on the auditory non-social (non-vocal) > auditory social (vocal) contrast (significant increase in HbO and/or significant decrease in HbR). A mask was applied such that we considered a significant activation on the contrast only if there was a significant activation to auditory non-social (non-vocal) compared to baseline in the first place. No channel showed a significant contrast.

